# Representation of Visual Landmarks in Retrosplenial Cortex

**DOI:** 10.1101/811430

**Authors:** Lukas F. Fischer, Raul Mojica Soto-Albors, Friederike Buck, Mark T. Harnett

**Affiliations:** Department of Brain & Cognitive Sciences and McGovern Institute for Brain Research, Massachusetts Institute of Technology, Cambridge, MA, 02139, USA

## Abstract

The process by which visual information is incorporated into the brain’s spatial framework to represent landmarks is poorly understood. Studies in humans and rodents suggest that retrosplenial cortex (RSC) plays a key role in these computations. We developed an RSC-dependent behavioral task in which head-fixed mice learned the spatial relationship between visual landmark cues and hidden reward locations. Two-photon imaging revealed that these cues served as dominant reference points for most task-active neurons and anchored the spatial code in RSC. Presenting the same environment but decoupled from mouse behavior degraded encoding fidelity. Analyzing visual and motor responses showed that landmark codes were the result of supralinear integration. Surprisingly, V1 axons recorded in RSC showed similar receptive fields. However, they were less modulated by task engagement, indicating that landmark representations in RSC are the result of local computations. Our data provide cellular- and network-level insight into how RSC represents landmarks.

## Introduction

Spatial navigation requires the constant integration of sensory information, motor feedback, and prior knowledge of the environment (Hardcastle, Ganguli, & Giocomo, 2015; McNaughton, Battaglia, Jensen, Moser, & Moser, 2006; Taube, 2007; Valerio & Taube, 2012). Visual landmarks are particularly advantageous for efficient navigation, representing information-rich reference points for self-location and route planning (Etienne, Maurer, & Séguinot, 1996; Gothard, Skaggs, & McNaughton, 1996; Jeffery, 1998; Knierim, Kudrimoti, & McNaughton, 1995; McNaughton, Chen, & Markus, 1991). Even in situations where the immediate surroundings may not be informative, distal landmarks can provide critical orientation cues to find goal locations (Morris, 1981; Tolman, 1948). Their importance is further underlined by the fact that salient visuo-spatial cues anchor almost every type of spatially-tuned cells observed in the mammalian brain to date, including head direction cells (Jacob et al., 2017; Taube, Muller, & Ranck, 1990a; Yoder et al., 2011), hippocampal place cells (Buzsáki, 2005; Jeffery, 1998), and grid cells in the medial entorhinal cortex (Hafting, Fyhn, Molden, Moser, & Moser, 2005; Pérez-Escobar, Kornienko, Latuske, Kohler, & Allen, 2016). Even in scenarios where self-motion feedback is in conflict with external visual cues, landmarks exert powerful influence on the head direction system (Etienne & Jeffery, 2004; Valerio & Taube, 2012). Additionally, a number of theoretical studies have shown the importance of landmarks for error correction during spatial computations (Burgess, Barry, & O’Keefe, 2007; Fuhs & Touretzky, 2006; Monaco, Knierim, & Zhang, 2011; Sreenivasan & Fiete, 2011). However, the process by which visual information is incorporated into the brain’s spatial framework remains poorly understood.

Converging evidence points to the retrosplenial cortex (RSC) as an important locus for landmark computations. Studies in human patients with damage to RSC as well as functional imaging studies suggest a key role for RSC in utilizing familiar visual cues for navigation (Auger, Mullally, & Maguire, 2012; Epstein, 2008; Epstein & Vass, 2014; Maguire, 2001; Vann, Aggleton, & Maguire, 2009). Additionally, RSC exhibits some of the earliest measurable metabolic dysfunction in Alzheimer’s disease (AD) (Minoshima et al., 1997; Pengas, Hodges, Watson, & Nestor, 2010; Villain et al., 2008). This is consistent with the putative roles of RSC in general mnemonic processing (Cooper, Manka, & Mizumori, 2001; Spiers & Maguire, 2006; Svoboda, McKinnon, & Levine, 2006; Valenstein et al., 1987) and route-planning (Spiers & Maguire, 2006), both of which are hallmarks of cognitive decline in AD patients (Vann et al., 2009). Lesion studies in rodents indicate that RSC is also important for navigating based on self-motion cues alone (Elduayen & Save, 2014). Together, these findings are congruent with known RSC anatomy: situated at the intersection of areas that encode visual information, motor feedback, higher-order decision making, and the hippocampal formation (Groen & Wyss, 1992; Kononenko & Witter, 2012; Miyashita & Rockland, 2007; Sugar, Witter, van Strien, & Cappaert, 2011), RSC is ideally positioned to integrate these inputs to guide ongoing behavior. Electrophysiological recordings in freely moving rats have shown that individual RSC neurons conjunctively encode space in egocentric and allocentric spatial reference frames (Alexander & Nitz, 2015). When placed in a one-dimensional environment, RSC neurons exhibit single, spatially tuned receptive fields (Mao, Kandler, McNaughton, & Bonin, 2017), while in two-dimensional environments (Alexander & Nitz, 2017) they were found to express multiple receptive fields. RSC neurons have further been shown to encode context as well as task-related cues such as goal location (Smith, Barredo, & Mizumori, 2012; Vedder, Miller, Harrison, & Smith, 2016). Recent studies have focused on understanding how accumulation of evidence (Koay, Thiberge, Brody, & Tank, 2019) and/or locomotion (Clancy, Orsolic, & Mrsic-Flogel, 2019) are differentially represented in RSC and visual cortex. Finally, a subset of cells in RSC encode head direction in a way that is particularly sensitive to local environmental cues (Jacob et al., 2017). A common theme across these studies is the importance of visual inputs for RSC function. While the role of proximal, non-visual cues, such as whisker stimulation, has not been thoroughly evaluated, it is clear that visual cues alone are sufficient to guide behavior. Together, these converging results strongly implicate RSC as an important neural substrate for landmark encoding.

We set out to identify how visual cues within a virtual environment that inform an animal about a goal location are represented in RSC. To this end, we developed a task in which animals are required to learn the spatial relationship between a salient visual cue in the virtual environment and a rewarded zone, therefore making the cue a landmark. Studies investigating how spatial tuning is influenced by the environment generally use a single orienting cue (Hafting et al., 2005; Muller & Kubie, 1987; Taube, Muller, & Ranck, 1990b), visual cues directly indicating the presence or absence of a reward (Pakan, Currie, Fischer, & Rochefort, 2018; Poort et al., 2015) or cue-rich environments where understanding the visual surrounding was not required to locate rewards (Campbell et al., 2018; Fiser et al., 2016; Gauthier & Tank, 2018; Harvey, Collman, Dombeck, & Tank, 2009; Saleem, Diamanti, Fournier, Harris, & Carandini, 2018). In contrast, our task requires mice to use allocentric inputs as reference points, and combine them with self-motion feedback to successfully execute trials. We found the majority of task-active neurons, as well as the population response, to be anchored by landmarks, though this differed between layers 2/3 and 5. Showing the same visual stimuli at a static flow speed while animals were not engaged in the task resulted in significantly degraded responses, suggesting that active navigation plays a crucial role in RSC. Further analysis showed that landmark responses were the result of supralinear integration of visual and motor components. To understand how visual information is translated into behaviorally relevant representations in RSC, we recorded the activity of axons from the primary visual cortex (V1) during task execution. V1 sends strong projections to RSC (Oh et al., 2014) which, in turn, sends a top-down projection back to V1 (Makino & Komiyama, 2015), creating a poorly understood cortico-cortical feedback loop between a primary sensory and associative cortex. Understanding this circuit could provide key insights into how sensory and contextual information combine to guide behavior. We found strikingly similar receptive fields as those expressed by RSC neurons, suggesting that V1 inputs may be key in shaping their receptive fields. Importantly, their activity was less modulated by active navigation, illuminating a key difference between primary sensory and associative cortex.

## Results

### An RSC-dependent visual landmark navigation task

We developed a behavioral task that requires mice to use landmarks to locate hidden rewards along a virtual linear corridor (Fig. 1A and 1B). Each trial began at a randomized distance (50 - 150 cm) from one of two salient visual cues with a vertical or diagonal stripe pattern respectively. Along the rest of the corridor, a gray-and-black dot uniform pattern provided optic flow feedback but no spatial information. An unmarked 20 cm wide reward zone was located at fixed distances (80 or 140 cm, respectively) from the visual landmarks. A trial ended when an animal either triggered a reward by licking within the reward zone or received a ‘default reward’ when it passed through the reward zone without licking (at 100 cm or 160 cm distance from the landmark). Default reward was present throughout the experiment but constituted only a small fraction of trials in trained animals (mean ± SEM: 14.24 ± 2.65 % of trials, n = 13 sessions, 7 mice). The visual appearance of the hallway was infinitely long so that animals could not use the end of the track as a spatial cue. This task design forced animals to use the visual cue as a spatial reference point that informs them about the location of the reward. At the end of each trial, the animal was ‘teleported’ into a ‘black box’ for at least 3 seconds. The 3 second black box timeout was included in the task to give the animals a salient signal for the end and start of trials. It further ensured that GCaMP6f signals underlying different aspects of behavior (reward delivery and consumption versus the initiation of a new trial and changes in locomotion) could be disambiguated. Mice learned to use the visual cues to locate the reward zones along the corridor (Fig. 1C, mean 32.3 ± 3.6 training sessions for n = 10 mice). In theory, animals could achieve a high fraction of successful trials by licking frequently but randomly, or in a uniform pattern. We tested if we could use licking as a behavioral assay for an animal’s understanding of the spatial relationship between visual cue and reward location by calculating a spatial modulation index using a bootstrap shuffle test (supp. Fig. 1E and 1F, methods).

**Figure 1:**
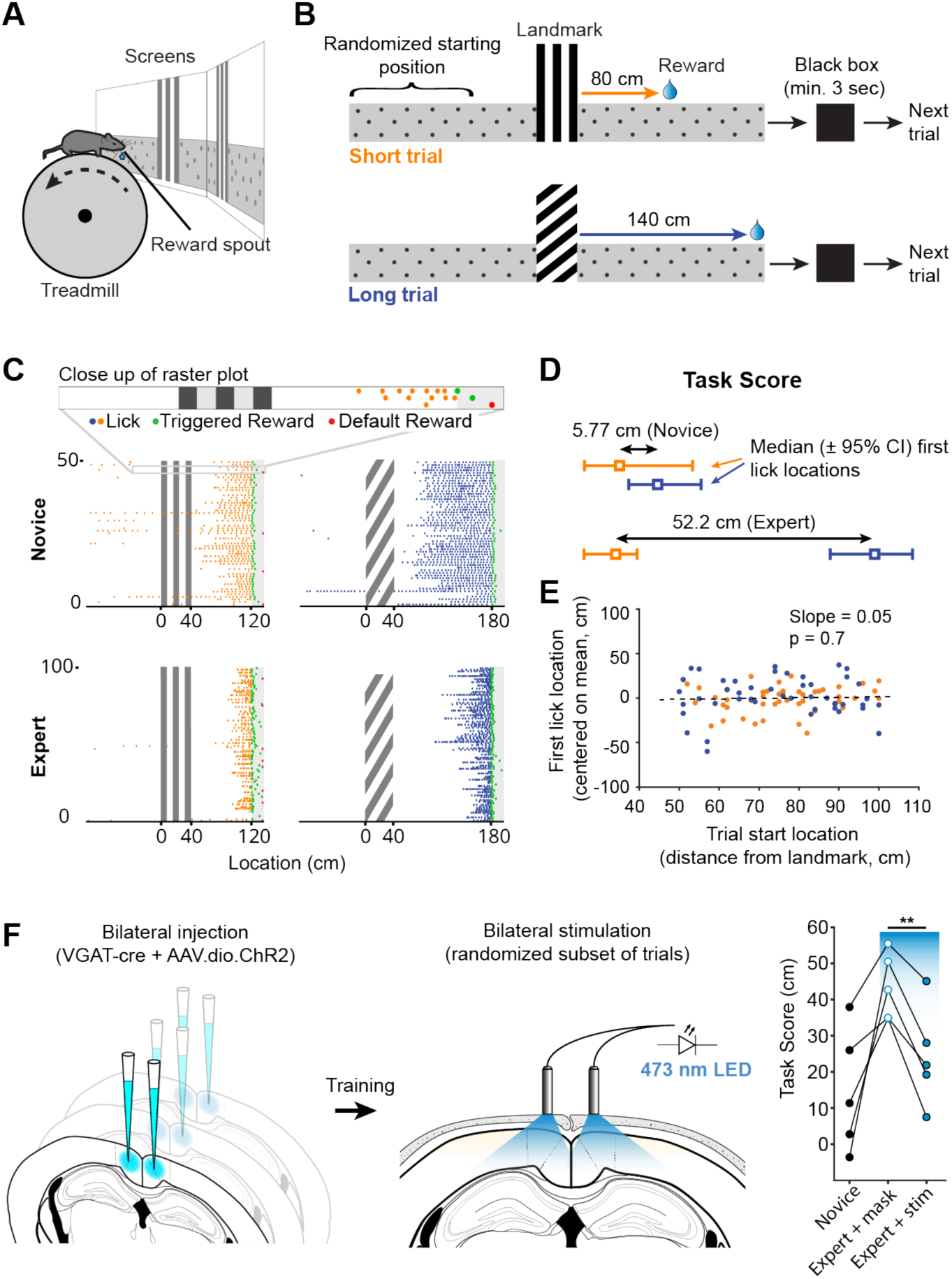
Landmark-dependent navigation task in virtual reality. (**A**) Schematic of experimental setup: mice are head-fixed atop a cylindrical treadmill with two computer screens covering most of the animal’s field of view. A reward spout with attached lick-sensor delivers rewards. (**B**) Task design. Animals learned to locate hidden reward zones at a fixed distance from one of two salient visual cues acting as landmarks. The two landmarks were interleaved within a session, either randomly or in blocks of 5. After each trial animals were placed in a ‘black box’ (screens turn black) for at least 3 seconds. The randomized starting location ranged from 50 – 150 cm before the landmark. (**C**) Licking behavior of the same animal at novice and expert stage. Expert animals (bottom) lick close to the reward zones once they have learned the spatial relationship between the visual cue and reward location. (**D**) How well mice were able to use landmarks to locate rewards was quantified by calculating the difference of the median location of licking onset for each landmark per session. (**E**) Relationship between trial start and first lick locations for one example session. Experimental design ensured that alternative strategies, such as using an internal odometer, could not be used to accurately find rewards. (**F**) RSC inactivation experiment. VGAT-Cre mice were injected with flexed Channelrhodopsin-2 (left). Stimulation light was delivered through skull-mounted ferrules on a random subset of trials (middle). During inactivation trials, task score was reduced significantly (right).

We then quantified an animal’s ability to use landmarks for navigation by calculating the difference between the median location of first licks on short and long trials, expressed as a “task score” (mean ± SEM of recording sessions: 34.92 ± 4.96 cm). Location of first licks as opposed to mean lick location or lick frequency was used as it provided the most conservative measure of where animals anticipated rewards. This task structure inherently minimizes the ability of mice using alternative strategies such as time or an internal odometer to locate rewards. We tested whether animals used the start of the trial and a fixed distance, time, or number of steps before they started probing for rewards (Fig. 1E). For each trial in one session, we plotted the location of the start vs. the location of the first lick and evaluated the linear regression coefficient showing no dependence of first lick on trial start location (mean ± SEM of slope: 0.14 ± 0.042, n = 13 sessions).

To test if RSC was involved in task performance, we bilaterally injected an AAV expressing a Cre-dependent channelrhodopsin-2 (ChR2) construct in multiple locations along the anterior-posterior axis of RSC (2-3 injections per hemisphere) in VGAT-Cre mice. This restricted ChR2 expression to GABAergic neurons in RSC and allowed us to rapidly and reversibly inhibit local neural activity (Lewis et al., 2015; Liu et al., 2014). Ferrules were implanted on the surface of the skull over RSC to deliver light during behavior. Stimulation was delivered by a 470 nm fiber-coupled LED on a randomized 50% subset of trials. The stimulation light was turned on at the beginning of a given trial and lasted until the end of the trial or the maximum pulse duration of 10 seconds was reached. A masking light was shown throughout the session. Task score on trials with stimulation was significantly lower compared to trials where only the masking light was shown within the same session (43.7 ± 7.41 cm vs. 24.3 ± 6.2 cm, n = 5 mice, paired t-test: p = 0.003, Fig. 1F), indicating that RSC activity contributes to successful execution of this behavior.

### Trial onset, landmark, and reward encoding neurons in RSC

We sought to understand which task features were represented by neurons in RSC. Mice injected with AAV expressing the genetically encoded calcium indicator GCaMP6f were trained until they reliably used landmarks to locate rewards. On average, we recorded from 121.0 ± 27.3 RSC neurons per mouse (n = 7 mice). GCaMP signals of all active neurons (> 0.5 transients/min, n = 1026) were tested for significant peaks of their mean response above a shuffled distribution (z-score of mean trace > 3, see methods) and for transients on at least 25% of trials. Neurons that met these criteria (n = 509) were then aligned to each of three points: trial onset, landmark, and reward (Fig. 2A, right; Fig. 2B). The peak activity of the mean GCaMP trace of each neuron with significant task modulation was then compared across alignment points and classified based on which task feature resulted in the largest mean response (Fig. 2B). The vast majority of neurons showed a single peak of activity in our task. Previous studies have found multiple peaks or sustained firing in RSC neurons (Alexander & Nitz, 2015, 2017). However, in contrast to these studies, our task did not contain repeating sections found in a W-shaped or plus-shaped maze. A peak response did not necessarily need to happen directly at a given alignment point, but could also occur at some distance from it. A landmark-aligned neuron, for example, did not have to exhibit its peak response at the landmark (Fig. 2F). This analysis was carried out for short trials and long trials independently. Our results show that the majority of RSC neurons found to be task engaged were aligned to the visual landmark (Fig. 2C, n = 61 short trial onset and 63 long trial onset, 217 and 253 landmark, 99 and 118 reward neurons, respectively; 7 mice; mean ± SEM fractions of aligned neurons: trial onset: 5.5 ± 0.1%, landmark: 22.2 ± 1.7%, reward: 11.5 ± 0.9%; one way ANOVA_short_: p = 0.0028; ANOVA_long_: p < 0.001, Tukey’s HSD post-hoc pairwise comparison with Bonferroni correction). A smaller but sizeable fraction of RSC neurons were aligned to the reward point, suggesting that RSC encodes behavioral goals as well. We further found neurons that reliably encoded the onset of a trial, regardless of where an animal was placed on the track relative to the visual landmark. Consistent with previous findings (Alexander & Nitz, 2015), these data indicate that egocentric (trial onset) as well as allocentric (landmark and reward) variables are encoded in RSC during landmark-based navigation. We employed a template matching decoder (Montijn, Vinck, & Pennartz, 2014) to analyze how well trial type could be decoded from neural activity alone (Fig. 2E). While trial onset neurons provided only chance level decoding (mean 45.6 ± 5.6% correct), trial type decoding by landmark neurons was significantly higher (77.4 ± 6.2%, Kruskal-Wallis test p = 0.0118; post-hoc Mann-Whitney U pairwise testing with Bonferroni correction for multiple comparisons). Decoding based on reward-aligned neurons was poor, but above chance (62.3 ± 5.6%), suggesting that landmark neurons carry most information about which type of trial the animal is currently executing.

**Figure 2:**
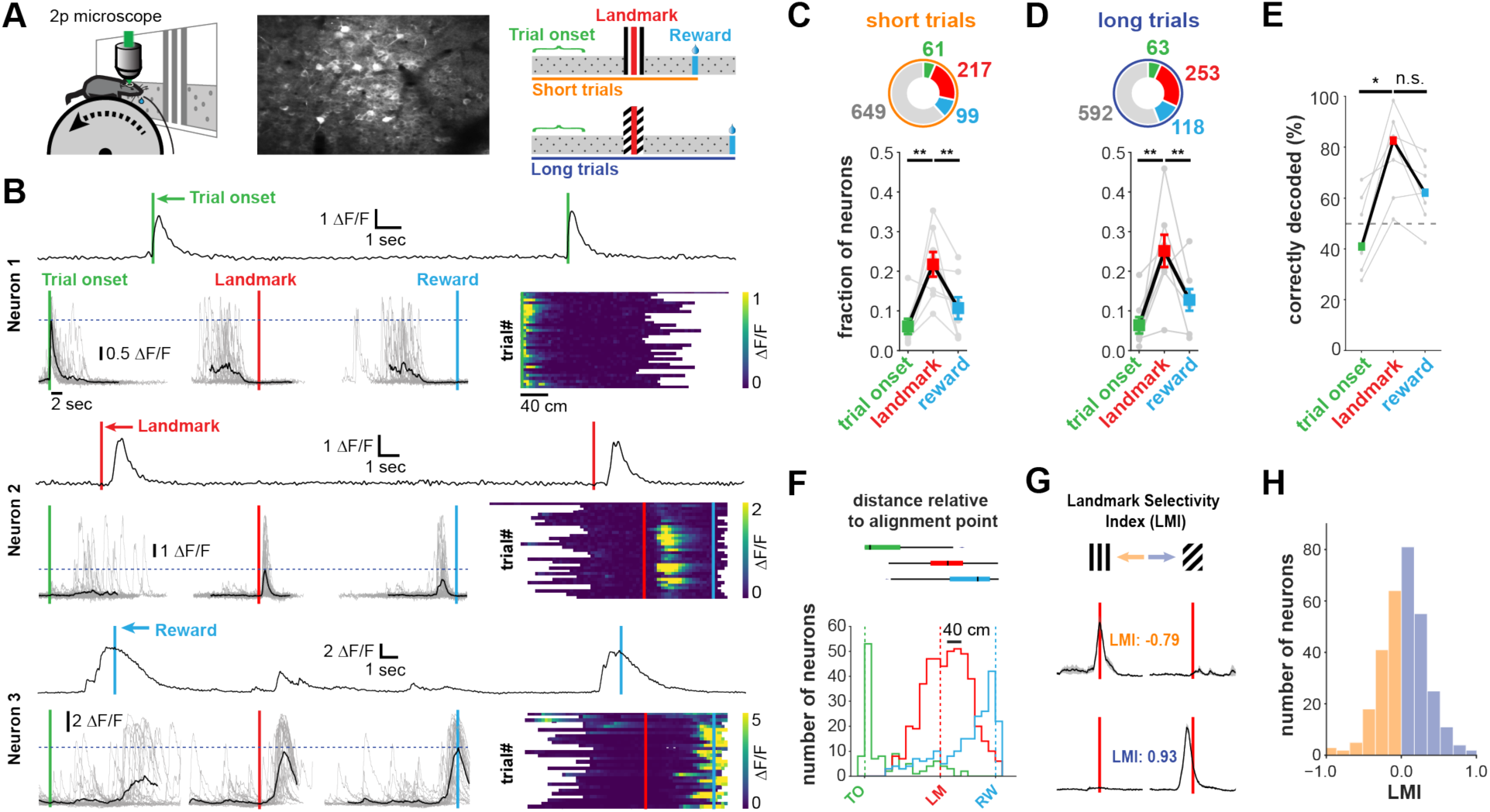
Neuronal responses in RSC during landmark-dependent navigation. (**A**) Overview: GCaMP6f-positive neurons were recorded with two-photon microscopy in expert animals executing the task. Activity of each neuron was then aligned to each of three points: trial onset (green), landmark (red), and reward (cyan) (independently for short and long trials). (**B**) Each neuron’s best alignment point was assessed by quantifying the peak of its mean trace and comparing it to the other alignment points. Rows show trial onset (top), landmark (middle), and reward aligned (bottom) example neurons. (**C, D**) Alignment of task-active neurons. The majority of task-engaged neurons were aligned to the landmark on short (C) and long (D) trials (n = 7 mice). (**E**) We applied a template matching decoder (Montijn et al., 2014) to decode the trial type based on the neural responses recorded from each animal. Trial onset neurons provided chance level decoding. However, landmark neurons provided significantly higher decoding accuracy which remained elevated for reward neurons (**F**) Mean distance of transient peaks of individual neurons relative to alignment point. Boxplot shows the spatial distribution (1^st^ - 3^rd^ quartile) of the histogram below. (**G**) Two landmark-selective neurons. Landmark selectivity was calculated as the normalized difference between peak mean responses. (**H**) The landmark selectivity index (LMI) of all landmark neurons shows a unimodal distribution.

We asked whether the subpopulation of landmark encoding neurons showed a preference for visual cue identity, in which case we expected a bimodal distribution. Alternatively, a unimodal distribution would provide evidence for the encoding of landmarks as an environmental feature informing the animal about a goal location. We calculated a landmark modulation index as the difference between peak activity divided by the sum of their activity [LMI = (LM_short_ – LM_long_)/(LM_short_ + LM_long_)]. Peak activity for each trial type was calculated separately such that shifts in relative distance to the landmark are taken into account. The distribution shows that most neurons do not show a specific preference for landmark identity, though there are a small number of neurons that are tuned to landmark identity (Fig. 2H). In contrast, trial onset neurons showed less trial type selectivity (supp. Fig. 5E). Together these results show that neurons encode a mix of task variables with a strong preference for visual cues informing the animal about goal locations.

### Landmarks anchor the representation of space in RSC

We tested how well an animal’s location is represented in RSC using population vector cross-correlation for all task active neurons (n = 509, K. M. Gothard, Skaggs, and McNaughton 1996; Alexander and Nitz 2015, 2017; Mao, Kandler, McNaughton, and Bonin 2017). The vectors were constructed by randomly drawing half of the trials for a given neuron to create one vector and the other half for the second vector. Activity was binned into 5 cm wide bins and the mean across all included trials was calculated and normalized to 1. In this task, trial start provided the animal with information on trial only fixed reference points to locate rewards along the virtual corridor. We therefore tested how well the population was anchored by trial start or landmarks by measuring local population vector cross-correlation aligned to those two points (Fig. 3A-F, left vs. right columns, G). If trial onset provided the dominant reference point for the representation of space, it would suggest that a mouse’s self-motion cues, such as number of steps since the start of a trial, drive responses in RSC. In contrast, landmark-anchored responses would indicate that allocentric cues, in combination with self-motion feedback, are the principal drivers of activity. We found largely even tiling of space from the trial onset until reward (Fig. 3A and 3D). To test the spatial specificity of the population code, we calculated a population vector cross-correlation matrix using the Pearson cross-correlation coefficient (Fig. 3B and 3E) for each spatial bin. To ensure that randomly splitting data into halves didn’t lead to spurious results, we calculated the mean of 100 cross-correlation maps, each randomly drawing a different subset of trials. Slices of the cross-correlation matrix (Fig. 3C and F, taken at the dashed lines indicated in 3B and 3E), reveal that the spatial code is sharpest at the landmark. The cross-correlation at the animal’s true location (i.e. along the diagonal from top left to bottom right) significantly increases as the animal approaches the landmark and remains elevated until it reaches the reward (Fig. 3G). We tested how well we could reconstruct the animal’s location from the neural code by calculating how far the pixel with the highest cross-correlation was from the actual location for each row in the cross-correlation matrix. We observed a significantly lower location reconstruction error when neural activity was aligned to landmarks, rather than trial onset (mean ± SEM: 3.26 ± 0.63 vs. 4.78 ± 0.43 short trials; 3.87 ± 0.42 vs. 5.93 ± 0.38, unpaired, 2-tailed t-test: p < 0.05 (short), p < 0.001 (long)). Finally, we analyzed how well we could reconstruct animal location based on the neural population activity on the short track when animals were on the long track and vice versa (supp. Fig. 5A and 5B). Consistent with our finding that most neurons are active on both trials (Fig. 2H, supp. Fig. 5C), we found a small, not significant increase in reconstruction error (supp. Fig. 5D). These results provide evidence for a spatial code in RSC that is strongly modulated by environmental cues to inform the animal about the location of its goal.

**Figure 3:**
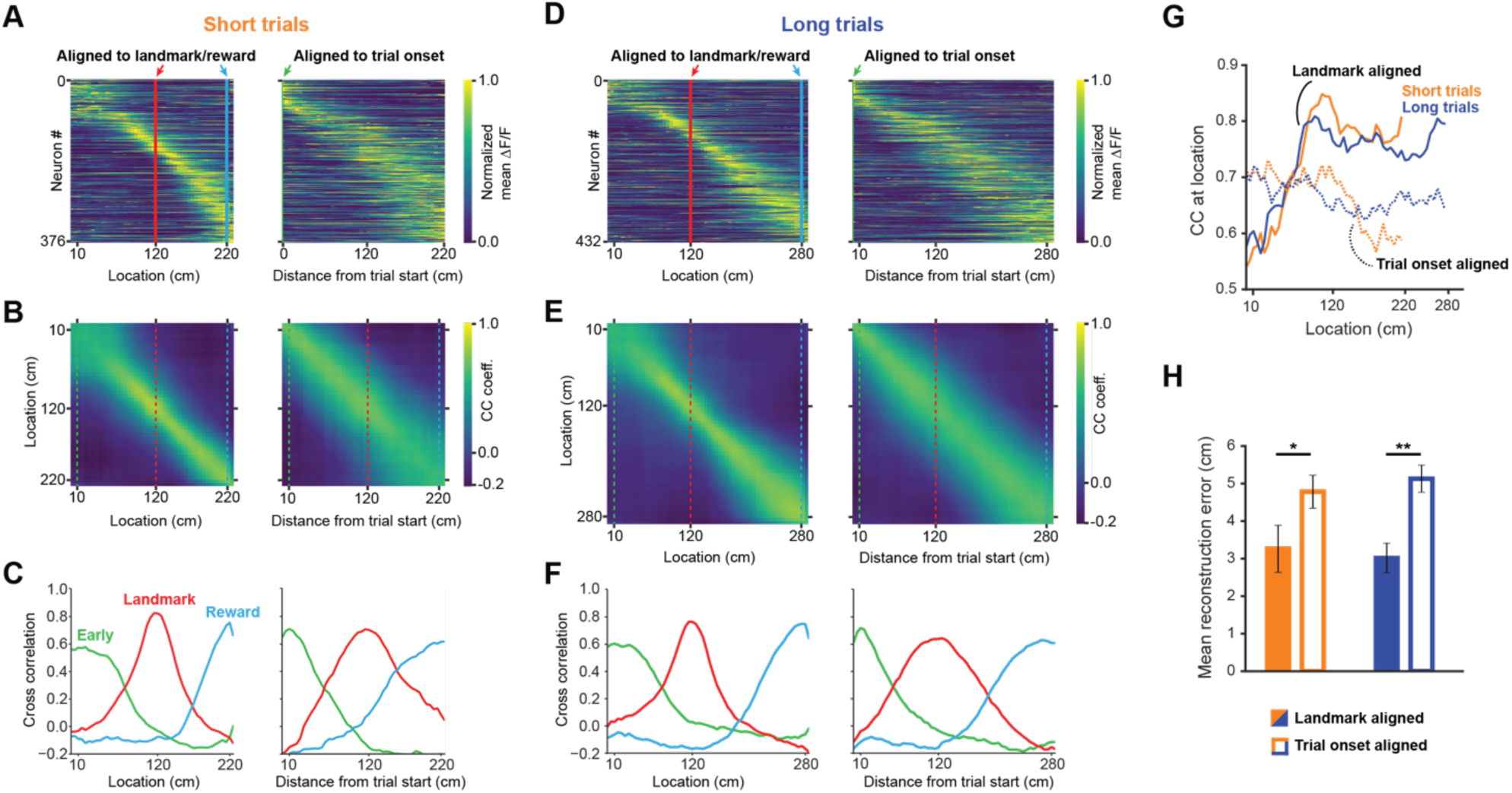
A landmark-anchored code for space in RSC. (**A**) Activity of all task engaged neurons ordered by location of peak activity on short trials. Left columns: neurons aligned to the landmark/reward; right columns: same neurons aligned to trial onset point. (**B**) Population vector cross-correlation matrices of data shown in (A). (**C**) Slices of the cross-correlation matrices early on the track (green dashed line), at the landmark (red dashed line), and at the reward point (blue dashed line), show sharpening of the spatial code at the landmark. (**D-F**) Same as (A-C) but for long trials. (**G**) Population vector cross-correlation values at the animal’s actual location. Solid lines: activity aligned to landmark/reward; dashed lines: activity aligned to trial onset. (**H**) Reconstruction error, calculated as the mean distance between the maximum correlation value in the cross-correlation matrices and the animal’s actual location, is significantly lower when neural activity is aligned to landmarks (solid bars) compared to trial onset aligned (open bars; unpaired t-test: short trials: p < 0.05, long trials: p < 0.001).

### Active task execution sharpens spatial tuning and increases robustness of responses in RSC

To determine if goal directed navigation, as opposed to visual input alone, was required to explain RSC activity, we recorded neurons while animals were shown the same virtual corridor without actively executing the behavior. During this decoupled stimulus presentation paradigm (DC), the virtual corridor moved past the animals at two speeds: 10 cm/sec and 30 cm/sec (Fig. 4A, see supp. Fig. 1 for speed profiles during virtual navigation). Both trial types were interleaved in the same way as during virtual navigation, but no rewards were dispensed when animals reached reward locations. We found a significant decrease in neuronal responses in this condition (Fig. 4B and 4C, mean ± SEM VR: 0.163 ± 0.004, mean decoupled: 0.0511 ± 0.004, paired, two-tailed t-test: p<0.0001). This was true for neurons of all three categories: trial onset, landmark, and reward (Fig. 4D, median values VR: trial onset = 0.1, landmark = 0.18, reward: 0.13; decoupled: trial onset = 0.02, landmark = 0.03, reward = 0.02). This result suggests that activity in RSC is strongly dependent on active task engagement. Congruent with this, population activity showed significantly less spatial specificity during decoupled stimulus presentation (Fig. 4E-H, mean reconstruction error ± SEM: 3.26 ± 0.62 vs. 7.44 ± 0.81 short trials; 3.02 ± 0.39 vs. 9.66 ± 0.96, unpaired, two-tailed t-test: p = 0.0001 (short), p < 0.0001 (long)). This indicates that encoding of behaviorally-relevant variables in RSC is dependent on ongoing behavior, rather than being driven solely by sensory inputs. Not providing a reward and decoupling the stimulus presentation from animal locomotion constitute simultaneous changes that may both influence neural activity. However, if reward anticipation was a key driver in the change in neural activity we would expect neurons anchored by trial onset or landmark to be less affected than reward driven neurons. We find that all neuron types are similarly affected (Fig. 4D), suggesting that reward anticipation is not the major cause for the change in activity we observed. A second potential factor modulating neuronal responses is whether the animal is attending to the cue or not. We have addressed this issue by analyzing responses during quiet wakefulness and locomotion in the next section (Fig. 5).

**Figure 4:**
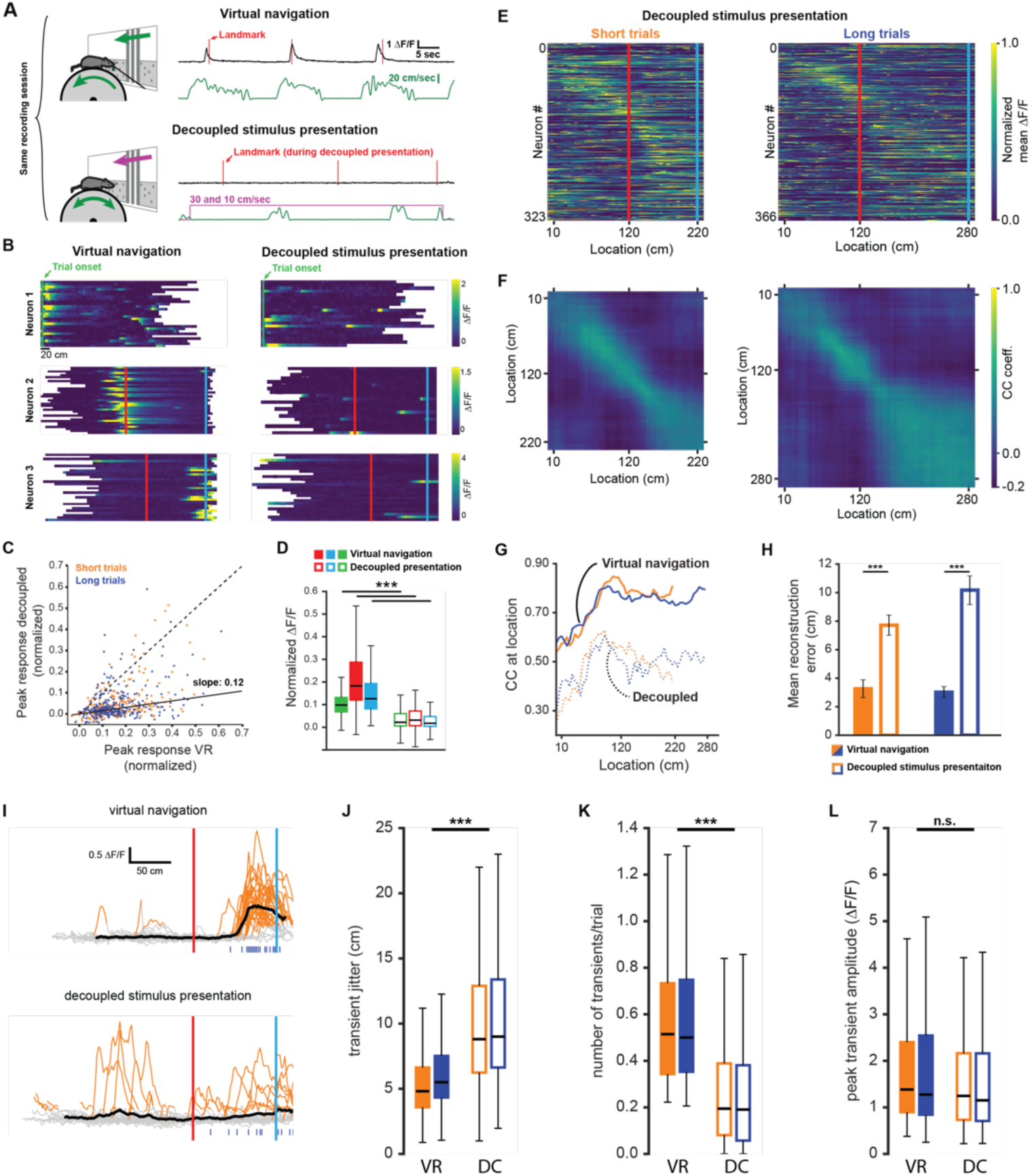
Neuronal activity during decoupled stimulus presentation. (**A**) Recording session structure: after recording from neurons during virtual navigation, the same stimuli were presented in an ‘open loop’ configuration where the flow speed of the virtual environment was decoupled from the animal’s movement on the treadmill. (**B**) Trial onset, landmark, and reward example neurons under these two conditions. (**C, D**) Response amplitudes of all task engaged RSC neurons during decoupled stimulus presentation (Kruskal-Wallis: p < 0.0001, Wilcoxon signed-rank pairwise comparison with Bonferroni correction indicated in (D)). (**E, F**) Population activity and population vector cross-correlation during decoupled stimulus presentation for short (left) and long (right) trials. (**G**) Local cross-correlation at animal’s location is smaller during decoupled stimulus presentation. (**H**) Mean location reconstruction error. Reconstructing animal location from population vectors is significantly less accurate when the animal is not actively navigating (unpaired t-test, short trials & long trials: p < 0.0001). (**I**) Traces of example neuron activity overlaid during virtual navigation (top) and decoupled stimulus presentation (bottom) with transients highlighted in orange. Ticks along bottom indicate peaks of transients around the neuron’s peak response. (**J**) Spread of transient peak location around peak mean response measured as the standard error of the mean of (standard error of mean of transient peak location – mean peak location). Solid bars: virtual navigation (VR), open bars: decoupled stimulus presentation (DC). (**K**) Average number of transients/trial during virtual navigation and decoupled stimulus presentation. (**L**) Average amplitude of transients in VR and DC conditions.

**Figure 5:**
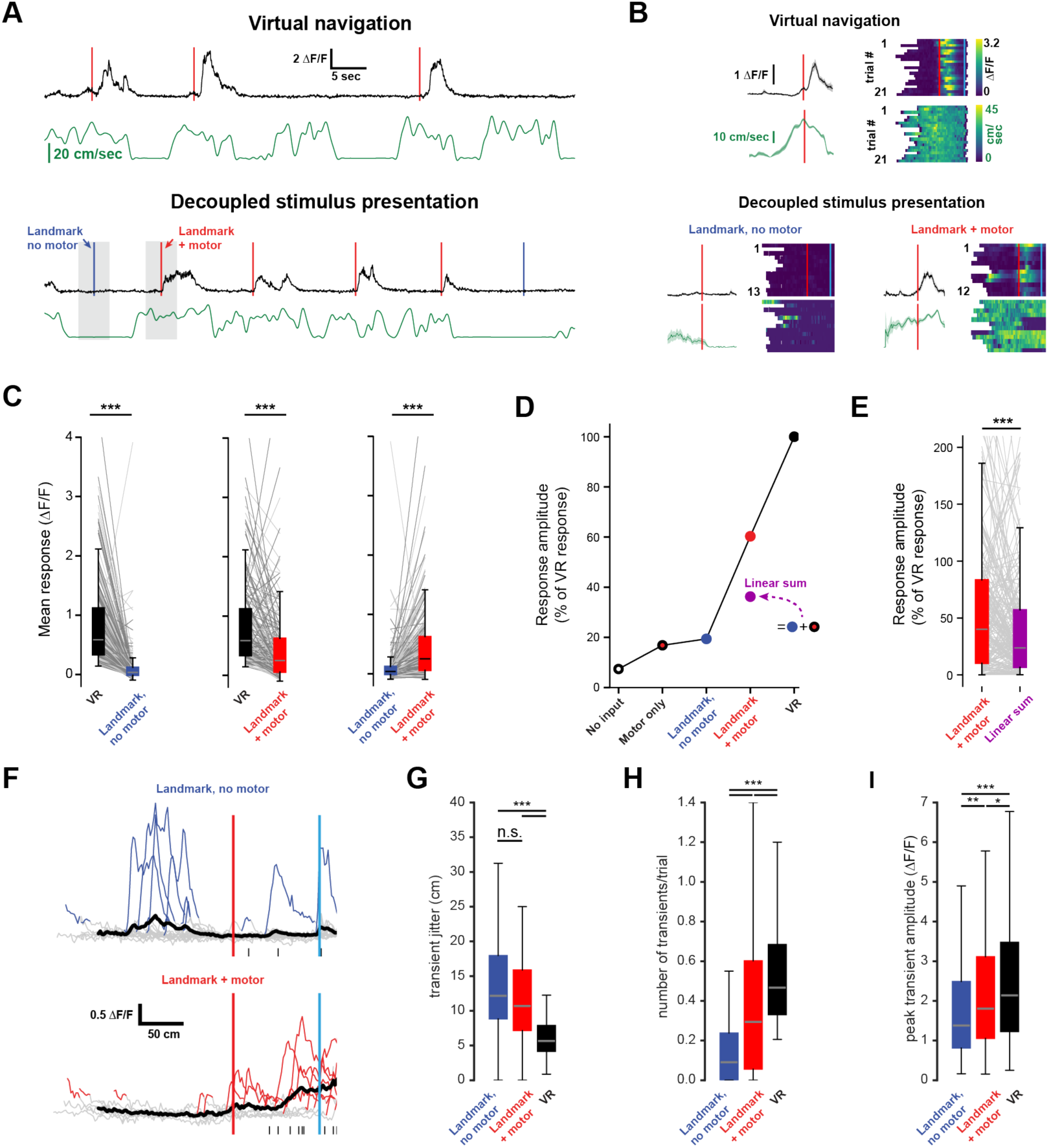
Non-linear integration of visual and motor inputs in RSC landmark neurons. (**A**) Example neuron during virtual navigation (top) and decoupled stimulus presentation as the animal is running or resting (bottom). (**B**) Same neuron as in (A) with all instances where the animal was running or resting, averaged (left) and raster plots of the whole session (right). (**C**) Activity of population during virtual navigation and decoupled stimulus presentation (**D**) Neuronal responses normalized to peak activity in VR under different conditions. ‘No input’ and ‘motor only’ responses were measured while animals are in the black box between trials (median and spread of data shown in (C)). (**E**) The sum of ‘landmark, no motor’ + ‘motor’ is smaller than ‘landmark + motor’ responses suggesting nonlinear combination of visual and motor inputs. (Wilcoxon signed-rank test, p < 0.0001). (**F**) Traces of example neuron when the animal is passively watching the scene (top) or locomoting (bottom). Blue ticks along bottom indicate transients around that neuron’s peak mean activity during virtual navigation (see Fig. 4I). (**G**) Spread of transient location around peak mean activity in VR (standard error of mean of transient peak location – mean peak location). (**H**) Average number of transients/trial and average amplitude (**I**) in both conditions. Kruskal-Wallis test p < 0.0001; Mann-Whitney-U pairwise comparisons with Bonferroni correction results indicated, * = <0.05, ** = <0.01, *** = <0.001.

We sought to gain further insight into potential mechanisms underlying the reduction in activity during decoupled stimulus presentation by comparing the patterns of GCaMP6f signals observed during virtual navigation and passive viewing. Individual events were detected when ΔF/F exceeded 6 standard deviations of a neuron’s baseline activity for at least 2 spatial bins (bin size: 2 cm) and lay within ±60 cm of the peak mean response (Fig. 4I). We found that the standard error of the distance of individual transients to the peak of the mean trace (Fig. 4J, median jitter (cm): short_VR_ = 4.81, short_DC_ = 8.8; long_VR_ = 5.5, long_DC_ = 9.0) was lower during virtual navigation compared to decoupled stimulus presentation. In other words, transients were more tightly clustered around that neuron’s peak response when the animal was actively engaged in the task. Furthermore, we saw a significant reduction in the number of transients per trial during decoupled stimulus presentation (Fig. 4K, median values (transients/trial): short_VR_ = 0.51, short_DC_ = 0.19; long_VR_ = 0.50, long_DC_ = 0.19), but only very little change in the amplitude of individual transients (Fig. 4L, median values (ΔF/F): short_VR_ = 1.38, short_DC_ = 1.25; long_VR_ = 1.27, long_DC_ = 1.15, Kruskal-Wallis test p < 0.0001; Mann-Whitney-U pairwise comparisons with Bonferroni correction results indicated, *** = <0.001). These results show that the reduced activity during decoupled stimulus presentation is due to poorer spatial anchoring of activity and fewer instances of a given neuron to exhibit a transient.

### Nonlinear integration of visual and motor inputs in RSC

Nonlinear integration of synaptic inputs dramatically enhances the computational power of individual neurons and neural networks (Miller and Cohen, 2001; London and Häusser, 2005; Mante *et al*., 2013; Rigotti *et al*., 2013; Jadi *et al*., 2014; Ranganathan *et al*., 2018). For a single neuron, the integration of multiple input streams may engage mechanisms of supralinear integration to produce complex, conjunctive responses (Bittner et al., 2015; Larkum, Kaiser, & Sakmann, 1999; Takahashi, Oertner, Hegemann, & Larkum, 2016; Xu et al., 2012). In contrast, neural networks may express conjunctive representations through high dimensional codes (Murray et al., 2017; Rigotti et al., 2013; Stringer, Pachitariu, Steinmetz, Carandini, & Harris, 2019). We therefore evaluated the evidence for nonlinear integration in landmark-anchored RSC neurons. During decoupled stimulus presentation, mice were free to spontaneously locomote on the treadmill or watch passively. We analyzed the activity of landmark neurons during periods when animals were locomoting (running speed > 3 cm/sec in a ±2 sec. window) versus when they were stationary (running speed < 3 cm/sec) as they passed through their receptive field, defined as the location of their peak mean activity during virtual navigation (Fig. 5A and 5B). We found that when landmark presentation occurred during locomotion, RSC landmark neuron activity was significantly increased (Fig. 5C, n = 203 neurons, 5 mice, mean ± SEM number of trials/neuron: 13.4 ± 0.36 resting, 8.8 ± 0.3 running; peak mean ΔF/F ± SEM: VR: 0.83 ± 0.05, landmark + motor: 0.47 ± 0.05, landmark, no motor: 0.12 ± 0.02, Kruskal-Wallis test: p = 4.9e-85, Mann-Whitney U post-hoc comparison with Bonferroni correction: p < 0.001 for all shown comparisons). However, we found that visual inputs alone or visual inputs plus locomotion did not elicit the same response as virtual navigation (Fig. 5D, Kruskal-Wallis test: p < 0.0001; post-hoc Mann-Whitney-U and Bonferroni correction: p < 0.0001 for all shown comparisons). We then evaluated the responses while animals were locomoting or stationary while in the ‘black box’ between trials to obtain estimates of population activity in ‘no input’ (neither visual nor motor inputs) and ‘motor only’ conditions (Fig. 5D, ΔF/F +/- SEM blackbox + motor: 0.14 ± 0.03, blackbox, no motor: −0.04 ± 0.02). Finally, we added ‘motor only’ and ‘landmark, no motor’ to calculate the linear sum of these inputs (Fig. 5D and 5E) and found that it was lower than ‘landmark + motor’. Analysis of transient patterns during ‘no motor’ and ‘+ motor’ conditions revealed that neurons show significantly more transients while locomoting (Fig. 5H, median values (transients/trial): no motor = 0.09, + motor = 0.29; VR = 0.46), however, transient amplitude and jitter were broadly similar (Fig. 5G and 5I, median jitter (cm): no motor = 12.17, + motor = 10.7; VR = 5.66; median amplitude (ΔF/F): no motor = 1.38, + motor = 1.8; VR = 2.14). We note that GCaMP6f may not provide linear translation from underlying spikes to fluorescence signal. However, our analysis focuses on relative differences within the same neuron under different conditions and thus nonlinearities of the calcium indicator are unlikely explain these results. This suggests that motor input drives RSC neurons, however it does not aid in anchoring their activity or modulating the number of spikes produced once it has reached firing threshold. Together these results provide evidence for substantial nonlinear integration of visual and motor inputs in RSC neurons during goal-directed virtual navigation as well as decreased, but still significant, nonlinear processing during decoupled stimulus presentation.

### Laminar differences in landmark encoding

To gain a deeper understanding of how cortical computations for landmarks may be implemented in RSC, we characterized the differences in the population code between layer 2/3 (L2/3) and layer 5 (L5) neurons. A previous study suggests that during linear locomotion on a belt, activity reminiscent of spatially tuned cells is broadly similar but sparser in deep compared to superficial layers (Mao et al., 2017). We examined how individual task features (landmark, trial onset, and rewards) are differentially represented in RSC L2/3 and L5. Cortical layers were identified by their depth under the dura, confirmed post-hoc with histological sections in a subset of animals. RSC does not contain a layer 4; L5 was separated from L2/3 by a small volume with low GCaMP-positive cell body density (Fig. 6A). Mean recording depth for L2/3 was 130.0 ± 4.0 µm and 327.0 ± 15.2 µm for L5 (likely corresponding to L5a). We found that superficial as well as deep layers contained trial onset, landmark, and reward neurons. However, L5 contained substantially fewer landmark neurons (Fig. 6B and 6C). Both layers were similarly modulated by virtual navigation, compared to decoupled stimulus presentation (Fig. 6D). These results are congruent with findings in Mao et al. 2017 and suggest that while both laminar microcircuits are similarly engaged, L2/3 may be key for encoding landmarks during ongoing behavior.

**Figure 6:**
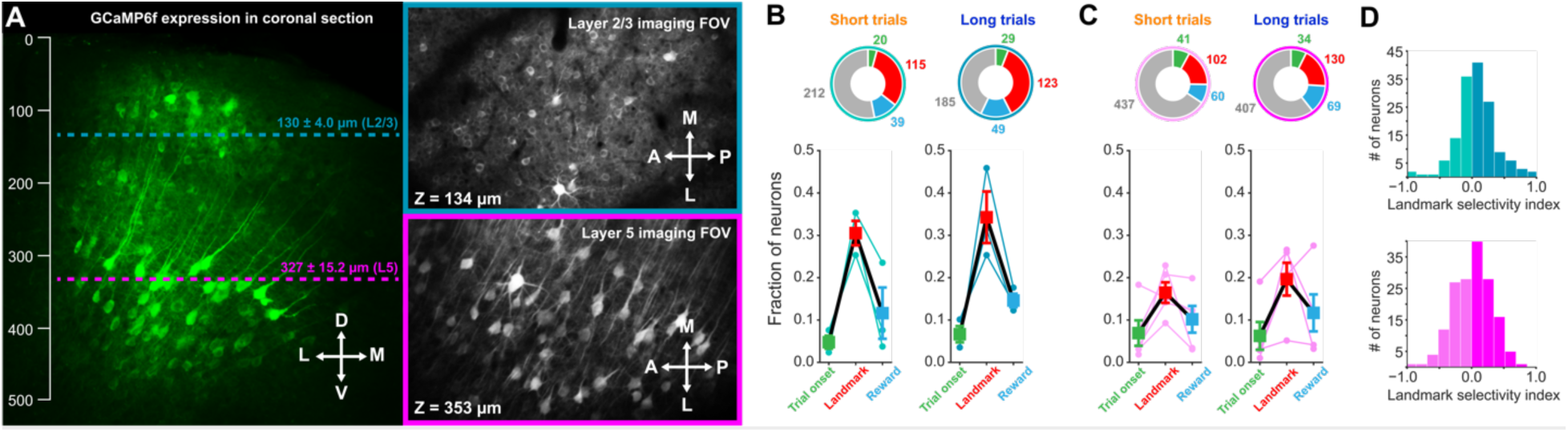
Differences in landmark encoding in layer 2/3 (turquois) and layer 5 (magenta). (**A**) Recording depths for imaging in superficial and deep layers indicated in a coronal section of GCaMP6f expressing neurons imaged post-hoc with confocal microscopy. Note the lack of layer 4 and that deep vs. superficial layers are separated by a region with few GCaMP-positive cell bodies. (**B, C**) Alignment of neuronal responses to task phases in superficial and deep layers separated by trial type. Layer 5 contained significantly fewer landmark-aligned neurons than layer 2/3 (One-way ANOVA, p < 10^-8^, Tukey HSD post-hoc test with Bonferroni correction). (**D**) Landmark selectivity is broadly similar between layers.

### V1 inputs to RSC represent task features but are less modulated by active navigation

Finally, we sought to dissect how RSC produces landmark representations by identifying what information it receives from primary visual cortex (V1), a major input source to RSC (Vogt & Miller, 1983). To this end, we injected GCaMP6f into V1 in a separate group of trained animals and recorded the responses of axonal boutons in RSC (Fig. 7A and 7B). Use of a passive pulse splitter (N. Ji, Magee, & Betzig, 2008) in the laser path allowed us to image axons continuously during self-paced behavioral sessions with no photobleaching or toxicity. To prevent overrepresentation of axons with multiple boutons in a given FOV, highly cross-correlated boutons were collapsed and represented as a single data point (see methods and supp. Fig. 4). In total, we found unique, task-related 77 axons in 4 animals. Unexpectedly, we found receptive fields that were strikingly similar to those we observed in RSC neurons (Fig. 7C and 7D). These boutons also tiled space along the virtual linear track in a parallel manner to RSC neurons (Fig. 7E and 7F). Furthermore, we found a similar preference of V1 boutons to be anchored by landmarks (Fig. 7G). However, when we quantified how active task engagement modulates activity in RSC neurons versus V1 boutons, we found that the former were significantly more modulated compared to the latter (Fig. 7 H and 7I, Mann-Whitney U: p < 0.001). These results are consistent with a number of recent studies that describe the encoding of non-visual stimuli in V1 (D. Ji & Wilson, 2007; Niell & Stryker, 2010; Pakan et al., 2018; Poort et al., 2015; Saleem et al., 2018). The specificity of these responses suggests that at least a subpopulation of RSC-projecting neurons in V1 is tuned to behaviorally-relevant visual cues to a previously unknown extent. Despite their specificity, however, they represent visual inputs more faithfully and are less modulated by context than RSC neurons themselves, pointing to the local computations performed in RSC.

**Figure 7:**
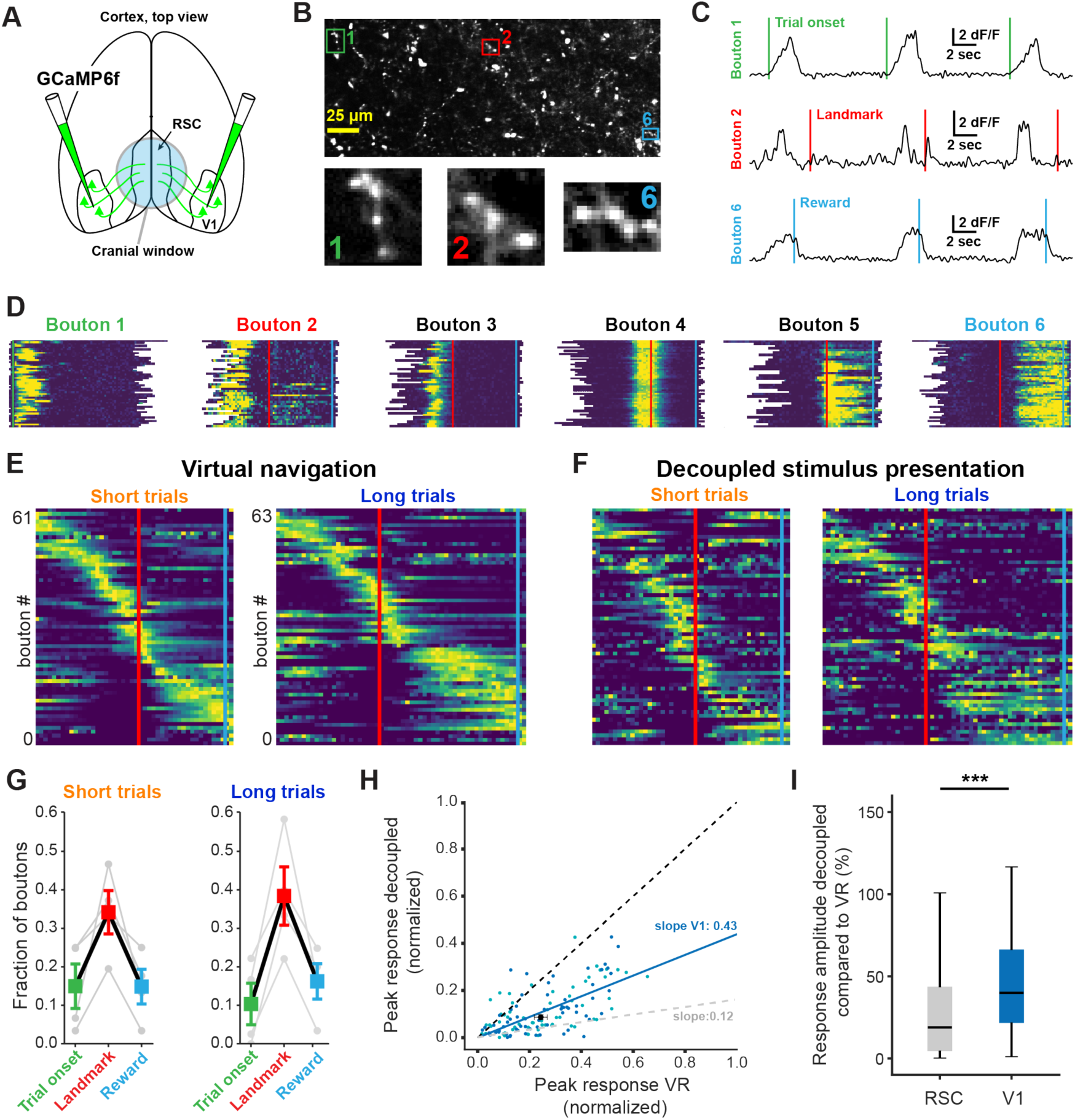
V1 axonal bouton responses in RSC. (**A**) Overview of injection and recording site. (**B**) Example FOV and three example boutons shown in (C) and (D). Where possible, ROIs were drawn around clusters of boutons belonging to the same axon. (**C**) Trial onset, landmark, and reward aligned boutons from same animal. (**D**) Six example boutons showing tuning to pre- and post-landmark portions of the track. (**E**) Population of V1 boutons in RSC ordered location of their response peak (n = 61 boutons short track/63 boutons long track, 4 mice). (**F**) Same boutons as in (E) during decoupled stimulus presentation. (**G**) Alignment of boutons to task features. (**H**) Response amplitude during virtual navigation of decoupled stimulus presentation with fitted regression line. In grey: fitted regression line for RSC neurons. (**I**) Comparison of response amplitude differences between VR and decoupled stimulus presentation in RSC neurons and V1 boutons (mean_RSC_ = 0.38 ± 0.03, mean_V1_ = 0.53 ± 0.04, Mann-Whitney U test: p < 0.001)

## Discussion

In this study, we introduce a novel behavioral paradigm in which mice learned the spatial relationships between salient environmental cues and goal locations (Fig. 1). The task required animals to discriminate visual cues, use them to localize themselves in space, and navigate to a rewarded zone based on self-motion feedback. Using this paradigm, we found that landmarks anchored the majority of neurons with task related responses (Fig. 2C and 2D) and significantly sharpened the representation of the animal’s current location in the population code (Fig. 3). These responses were not the result of simple visual and/or motor drive: showing the same visual stimuli while the animals were not engaged in the task elicited significantly attenuated responses (Fig. 4). Further dissection of neuronal activity provided evidence for supralinear integration of visual and motor information in RSC. Coinciding visual input and motor feedback during decoupled stimulus presentation did not elicit the same response amplitudes as observed during active navigation (Fig. 5). Interestingly, we found receptive fields expressed by V1 axonal boutons in animals executing the same behavior that were strikingly similar to those recorded from RSC neurons (Fig. 7). However, they were less modulated by active task engagement (Fig. 7H and 7I), indicating a hierarchy of sequential processing.

A major challenge in understanding how computations in RSC contribute to behavior is the diversity and complexity of functions attributed to this area (Maguire, 2001; Vann et al., 2009). Studies in humans suggest that RSC is key for utilizing environmental cues during navigation (Epstein, 2008; Ino et al., 2007; Maguire, 2001), while experiments in rodents found deficits in path integration (navigation based on self-motion cues) when RSC was lesioned or inactivated (Cooper et al., 2001; Cooper & Mizumori, 1999; Elduayen & Save, 2014). We describe a task that combines both of these navigational strategies: animals are required to use visual landmarks for self-localization, followed by path integration to successfully find rewards. Using optogenetic inactivation on a randomized subset of trials we found a deficit in the animal’s ability to use landmarks to guide localizing rewards (Fig. 1F). This could potentially be explained by a pure path integration deficit. However, if this was the case, RSC should exhibit a purely ego-centric representation of space, i.e. aligned to the start of a given trial. In contrast, we find that spatial representations in RSC are anchored by allocentric (landmark) cues and maintained by self-motion feedback after the animal has passed the visual cue (Fig. 3).

Numerous experimental and theoretical studies have emphasized the importance of landmarks for anchoring spatially tuned cells during navigation (Burak & Fiete, 2009; Campbell et al., 2018; Funamizu, Kuhn, & Doya, 2016; Gothard & Skaggs, 1996; Harvey, Coen, & Tank, 2012; Jeffery, 1998; Knierim, Kudrimoti, & McNaughton, 1998; Muller & Kubie, 1987), yet the mechanisms that combine inputs from different modalities to represent landmarks remain poorly understood. We found that simple linear summation of visual and motor inputs was insufficient to explain landmark encoding in RSC. Instead, a nonlinear mechanism, or multiple mechanisms, underlie the integration of these variables to produce robust visuo-spatial responses during navigation. Active navigation (in virtual reality) sharpened spatial tuning and increased robustness of encoding in RSC compared to viewing the movie without being engaged in the task. Interestingly, the amplitude of recorded transients was unchanged, suggesting the presence of a thresholding process in the circuit (Fig. 4I-L). Locomotion during decoupled stimulus presentation significantly increased the robustness of encoding while having very little effect on the fidelity of spatial tuning or transient amplitude. This indicates that motor input broadly pushes neurons towards spiking but does not contribute to its spatial tuning (Fig. 5F-I). Furthermore, our data shows that visual input alone is insufficient to explain the fidelity of spatial tuning we observed during virtual navigation (Fig. 5D). Our results indicate that the most likely mechanism underlying supralinear integration in RSC is the result of the multiplicative effects of significantly improved fidelity of spatial tuning and increased robustness of emitting transients within a neurons receptive field. While the latter seems to be mediated by motor inputs, the nature and source of an anchoring signal is unclear but may originate in the hippocampal formation where landmarks have been found to sharpen spatial tuning of neurons (Campbell et al., 2018; Gothard & Skaggs, 1996; Knierim et al., 1995).

The spatial tuning we observed in task active neurons in RSC appears similar to those of place cells (see Fig. 2B and Fig. 3A and 3D). This is consistent with findings in Mao et.al. 2017, who report spatial tuning in RSC that is somewhat degraded when tactile/visual cues are removed from a belted treadmill. However, RSC may not exhibit spatial representations that differ from CA1 when sensory information regarding goals is absent. Consistent with this, we find that RSC is strongly biased to encode behaviorally relevant visual cues that inform the animal about the location of a reward. These findings are complementary to previous studies showing that RSC conjunctively encodes information in egocentric and allocentric reference frames (Alexander & Nitz, 2015, 2017) as well as other variables (Smith et al., 2012; Vedder et al., 2016). However, in contrast to neurons expressing multiple receptive fields, as reported in Alexander and Nitz, 2015 and 2017, we found that most cells only had a single, stable receptive field. A possible explanation for this divergence is the design of our virtual environment, which did not have multiple discrete sections separated by turns.

RSC’s bias to encode behaviorally relevant stimuli is particularly interesting in light of its relationship with axonal inputs from V1 (Fig. 7). Using the same landmark-dependent navigation task, we found that V1 axons exhibited comparable receptive field tunings as RSC neurons. However, these responses were substantially less modulated by task engagement (Fig. 7I), suggesting that V1 axons encode visual features more faithfully. The modulation we did observe may be the result of strong top down inputs from RSC (Makino & Komiyama, 2015) or from other regions (Zhang et al., 2014). This is congruent with a recent study showing that activity in RSC is more correlated with V1 during locomotion compared to quiescent periods (Clancy et al., 2019). Indeed, before learning the behavioral significance of visual features in a novel environment, RSC may initially receive purely visual inputs from V1. As the animal learns to navigate in the new environment, feedback from RSC to V1 (as well as other areas such as ACC) may lead to modulated responses based on behavioral relevance in primary visual cortex, as reported by an increasing number of studies (Attinger, Wang, & Keller, 2017; Pakan et al., 2018; Poort et al., 2015; Saleem et al., 2018). Our data provides evidence that RSC may act as a critical processing node that gates behaviorally relevant visual inputs and relays them to the entorhinal cortex and other areas involved in spatial navigation, where its readout may be used to anchor spatially tuned neurons such as grid cells (Burak & Fiete, 2009; Campbell et al., 2018) or head-direction cells (Jacob et al., 2017). While RSC receives inputs from V2, M2, and other cortical areas (Oh et al., 2014; Sugar et al., 2011), functional imaging studies show RSC to be uniquely engaged during landmark based navigation (Auger et al., 2012; Epstein, 2008; Epstein & Vass, 2014; Maguire, 2001; Vann et al., 2009), suggesting RSC is indeed a key locus for integrating visual and spatial information compared to other association areas.

Finally, this work provides novel insights into the neural mechanisms underlying cognitive computations. Our results are consistent with data from humans with RSC lesions who show an impaired ability to use environmental cues for navigation, as well as neuroimaging studies that show increased activity in RSC during spatial behaviors (Cho & Sharp, 2001; Ino et al., 2007; Julian, Keinath, Marchette, & Epstein, 2018; Maguire, 2001; Robertson, Hermann, Mynick, Kravitz, & Kanwisher, 2016; Vann et al., 2009). Leveraging RSC to unravel how multiple input streams are integrated during higher level associative processes like navigation may in the future provide novel insights into the mechanisms of cognition and its dysfunction in Alzheimer’s disease and other currently intractable brain disorders.

## Methods

### 5.1 Animals and surgeries

All animal procedures were carried out in accordance with NIH and Massachusetts Institute of Technology Committee on Animal care guidelines. Male and female mice were singly housed on a 12/12 h (lights on at 7 am) cycle.

C57BL/6 mice were implanted with a cranial window and headpost at 7-10 weeks of age. First, the dorsal surface of the skull was exposed and cleaned of residual connective tissue. This was followed by a 3 mm wide round craniotomy centered approximately 2.5 mm caudal of the bregma. To minimize bleeding, particularly from the central sinus, the skull was thinned along the midline until it could be removed in two pieces. AAV1.Syn.GCaMP6f.WPRE.SV40 was injected at 2-6 injection sites, 350-600 µm lateral of the midline in boluses of 50-100 nl at a slow injection rate (max. 50 nl/min) to prevent tissue damage. Following injections, a cranial window was placed over the craniotomy and fixed with cyanoacrylate glue (Krazy Glue, High Point, NC, USA). The windows consisted of two 3 mm diameter windows and one 5 mm diameter window stacked on top of each other (Warner instruments CS-3R and CS-5R, Hamden, CT, USA). The windows were glued together with optical glue (Norland Optical Adhesive #71, Edmund Optics, Barrington, NJ, USA). Cranial windows consisted of 3 (instead of 2) stacked windows to account for increased bone thickness around the midline and minimize brain motion during behavior. Subsequently, the headplate was attached using cyanoacrylate glue and Meatbond® (Parkell Inc. NY, USA) mixed with black ink to avoid light leaking into the objective during recordings.

Mice prepared for imaging of V1 boutons in RSC had GCaMP6f injected into V1 (∼2.49 mm lateral, 3.57 caudal) through small burr holes at a depth of 600 µm to target primarily layer 5 neurons. For the rest of the procedure, the same steps as for imaging of RSC neurons were followed.

For the optogenetic inactivation during behavior experiment, VGAT-Ires-Cre mice knock-in on a C57BL/6 background (The Jackson Laboratory) were injected with flexed channelrhodopsin-2 (ChR2, AAV5.ef1a.DIO.ChR2.eYFP, University of Pennsylvania Vector Core) in 2 – 3 locations along the AP axis of RSC (50 – 100 nl per injection). Prior to injection the location of the central sinus was identified by placing saline on the skull and waiting until it was translucent. This was done because the overlying sagittal suture can be inaccurate in identifying the midline of the brain. One ferrule was placed centrally on each hemisphere over RSC. Each ferrule was calibrated prior to implantation to ensure the same light intensity was provided into each hemisphere.

### Virtual Reality Setup

Head-fixed mice were trained to run down a virtual linear corridor by locomoting on a polystyrene cylinder measuring 8 cm in width and 20 cm in diameter (Graham Sweet Studios, Cardiff, UK). The cylinder was attached to a stainless-steel axle mounted on a low-friction ball bearing (McMaster-Carr #8828T112, Princeton, NJ, USA). Angular displacement of the treadmill was recorded with an optical encoder (US Digital E6-2500, Vancouver, WA, USA). A custom designed head-restraint system was placed such that animals were comfortably located on the apex of the treadmill. Rewards were provided through a lick spout (Harvard Apparatus #598636) placed within reaching distance of the mice’s mouth. Timing and amount were controlled using a pinch valve (NResearch 225PNC1-21, West Caldwell, NJ, USA). Licking behavior was recorded using a capacitive touch sensor (SparkFun #AT42QT1010, CO, USA) connected to the lick spout. The virtual environment was created and rendered in real time in Matlab® using the software package ViRMeN (Aronov & Tank, 2014) as well as custom written code. Two 23.8” computer screens (U2414H, Dell, TX, USA) were placed in a wedge configuration to cover the majority of the mice’s field of view.

### Behavioral task design and training

After mice had undergone preparatory surgery, they were given at least one week to recover before water scheduling began. Initially, mice received 3 ml of water per day in the form of 3 g of HydroGel® (ClearH_2_O, Watertown, MA, USA), which was gradually reduced to 1.0-1.5 g per day. During this period, mice were handled by experimenters and habituated to being head restrained as well as running on a cue-less version of the virtual corridor. During habituation, mice were given small water rewards to allow them to acclimate to receiving 10% sugar-water rewards through a spout during head-restraint. Behavioral training began once mice were locomoting comfortably, as assessed by posture and gait. Initially, mice were trained on one trial type alone (short track). Each trial started at a randomized distance from the landmark (150-50 cm, drawn from a uniform distribution). The wall pattern consisted of a uniform pattern of black dots against a dark grey background to provide generic optic flow information. The view-distance down the corridor was not limited.

The landmark cues were 40 cm wide and extended above the walls of the corridor (see Fig. 1B). After passing the landmark, mice were able to trigger rewards by licking a fixed distance from the landmark. The reward zone was 20 cm long but not indicated in any way so that the animals had to use self-motion cues and the location of the landmark to locate it. If an animal passed through the reward zone without licking, an automatic ‘reminder’ reward was dispensed. Each reward bolus consisted of 4 - 6 µl of 10% sucrose water. Sucrose was added to maximize training success (Guo et al., 2014). Reward delivery marked the end of a trial and animals were “teleported” into a ‘black box’ for at least 3 seconds. In some training and recording sessions, animals were required to not lick or run for 3 seconds, however that requirement was later removed. Training using only one trial type was carried out daily in 30 to 60 minute sessions until licking behavior was reliably constrained to after the landmark. At that point, the second trial type (long track) was introduced. Training using two tracks was carried out until the licking behavior of mice indicated that they used landmark information to locate the reward (‘experts’, typically 2-4 weeks). An empirical bootstrap shuffle test was used to calculate confidence intervals and evaluate whether or not mean first lick locations where significantly different. At that point, mice were transferred to the 2-photon imaging rig. In some instances, a small number of training sessions with the recording hardware running were carried out on the imaging setup to acclimatize animals.

The spatial modulation z-score (SMZ) was calculated by randomly rotating the location of licks within each trial by a random amount. The fraction of correctly triggered trials within this new, shuffled session was calculated by evaluating whether at least one lick was within the rewarded zone. This process was repeated 1000 times and a null distribution of fraction successful from random licking was calculated.

### Optogenetic Inactivation Experiment

Optogenetic inactivation was carried out on mice that had been trained to expert level. Once they reached proficiency at using landmarks to locate rewards, the masking light was introduced. Animals were allowed a small number of sessions to habituate to the masking light before inactivation trials were introduced. The masking stimulus was provided by two 465 nm wavelength LEDs mounted on top of the computer screens facing the animal (Thorlabs LED465E, Thorlabs, NJ, USA). Optogenetic stimulation light was provided by a 470 nm fiber coupled LED (Thorlabs M470F3) powered by a Cyclops LED driver (Newman et al., 2015). Stimulation consisted of a solid light pulse with a maximum duration of 10 seconds (Lewis et al., 2015). Stimulation was provided on half of the trials in a random order, with the only exception that no two consecutive trials could be stimulation trials. Light intensity ranged from 1 - 10 mW and was calibrated individually for each animal. Each animal was observed during stimulation trials and checked for no visible effects on behavior such as change in posture or gait. No difference was found in mean running speed or licks per trial when the stimulation light on compared to when only the masking stimulus was shown (supp. Fig. 1). The task scores on mask only trials were compared to the task scores on mask + stimulation trials to assess deficits in the mice’s ability to use the landmark as a cue to locate rewards.

### Two-photon imaging

A Neurolabware 2-photon microscope coupled to a SpectraPhysics® Insight DeepSee II were used for GCaMP6f imaging. To prevent photodamage or bleaching during extended recording periods, a 4x pulse splitter was placed in the light path (N. Ji et al., 2008). The virtual reality software ran on a separate computer that was connected to the image acquisition system. Start and end of recording sessions were controlled by the virtual reality software to ensure synchrony of behavior and imaging data. Animals were placed in the head restraint and had a custom-designed 3D printed opaque sleeve placed over their cranial window to block light from the VR screens from leaking into the objective. The scope was lowered and suitable FOV identified before recordings began. Neurons in RSC were recorded at a wavelength of 980 nm. During V1 bouton recordings, the wavelength was switched to 920 nm. This was done to minimize autofluorescence from the dura mater, which is more pronounced at 980 nm excitation, especially during superficial recordings. Images were acquired at a rate of either 15.5 Hz or 31 Hz. In a subset of recordings, an electronically tunable lens was used to record from multiple FOVs in the same animal and session. In all but one cases, dual-plane imaging was used at a rate of 31 Hz, resulting in 15.5 Hz per plane acquisition. In a single recording session, 6 planes were acquired at 5.1 Hz. The two planes with most somas where included in this study. Recordings were acquired continuously throughout each session as opposed to epoch-based on trials.

### Image processing, segmentation, and signal extraction

Custom written Mathworks Matlab code was used for image registration, segmentation and signal extraction. Each recording session was stabilized using an FFT based rigid algorithm to register each frame to a template created from a subset of frames drawn at random from the whole session. This was followed by creating a pixel-by-pixel local cross-correlation and global cross-correlation maps. Regions of interest were drawn semi-automatically based on local cross-correlation from an experimenter defined seed-point. In addition to cross-correlation, global PCA, mean intensity, and other maps were created to aid identification of neurons and axonal boutons. Since the FOV was the same for virtual navigation and decoupled stimulus presentation, the same ROI maps created during virtual navigation could be used for decoupled stimulus presentation. During signal extraction, the mean brightness value of all pixels within a single ROI was calculated. A neuropil ‘donut’ was automatically generated around each ROI to allow for subtraction of local brightness from ROI signal. ΔF/F was calculated using a 60 second sliding time window. For neurons, F_0_ was calculated from the bottom 5^th^ percentile of data points within the sliding window. For boutons, the bottom 50^th^ percentile was used to calculate F_0_. Neuropil signal was subtracted from ROI signal prior to calculating ΔF/F. Each ROI time course was manually inspected prior to inclusion into analysis. ROIs were excluded if they had few transients (< 0.5/min). Transients were identified as detected whenever the ΔF/F signal was above 6 standard deviations for at least 500 ms. The ROI time course was then aligned and re-sampled to match behavioral data frame-by-frame using the Scipy signal processing toolbox (Jones, Oliphant, & Peterson, 2001). To test for long term imaging side effects despite using a pulse splitter, we tested for baseline drift of mean frame brightness for each included recording session (supp. Fig. 2).

### Neuron and axonal bouton classification

The time course of each neuron was split into individual trials and aligned to one of three anchor points: trial onset, landmark, and reward. For the neuron to be considered task engaged it had to fulfill the following criteria: 1) ΔF/F had to exceed 3 standard deviations of the ROI’s overall activity on at least 25% of all trials; 2) the mean ΔF/F across trials had to exceed a peak z-score of 3 at its peak. The z-score for each ROI was determined by randomly rotating its ΔF/F time course with respect to its behavior 500 times and the peak value of the mean trace was then used to calculate the peak z-score; 3) the minimum of the mean trace amplitude (i.e. highest – lowest value) had to exceed 0.2 ΔF/F. Criteria for axonal boutons were the same with the exception that the minimum mean trace amplitude was 0.1 ΔF/F. For the neurons that passed these criteria, the alignment point that resulted in the largest mean response was determined. To avoid edge-cases at the beginning and end of the track, the mean trace was only calculated for bins where at least 50% of trials were present. For the comparison of peak amplitude in Fig. 4C and 4D, the peak amplitude as a function of space, rather than time, was used. The landmark selectivity index was calculated for all neurons that were classified as landmark aligned on at least one trial type as LMI = *(LM_short_ – LM_long_)/(LM_short_ + LM_long_),* where LM_x_ refers to the peak response to the respective landmark. Only neurons that were classified as landmark-aligned neurons were included in that analysis. The fraction of neurons classified as trial onset, landmark, or reward were calculated from the total number of neurons with a baseline activity of at least 0.5 transients/min.

### Template matching decoder

A template matching decoder was used to assess the accuracy by which the trial type could be identified based on the activity of the different categories of neurons (trial onset, landmark, reward). First, template vectors were constructed for each trial type by calculating the mean response across trials within a session. The responses of the same neurons in individual trials were then compared to the template vectors, resulting in a similarity index for each trial type:

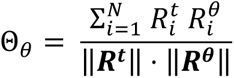

Here, Θ is the similarity index for trial type θ (short or long). ***R***^θ^ is the template vector for trial type θ, ***R^t^*** is a vector of the responses of all neurons in a given trial, and i…N are the indices of all neurons of a given category. Whichever similarity index was higher for a given trial was considered the decoded trial type and compared to the trial type the animal was actually on.

### Population plot and population vector analysis

Population plots were created by binning the activity of each neuron as a function of space. Each bin was 5 cm wide and all data points falling within a bin were averaged to calculate the mean activity at that location. The first bin started at 100 cm distance from the landmark such that it contained data from at least 50% of trials on average. The activity in each bin was normalized to the bin with peak activity of the same neuron such that all data ranged from 0 to 1. To plot the mean activity of all neurons in this study, the data was split. Half the trials were randomly drawn to calculate the bin with peak activity. The other half of the trials was used to calculate activity to be plotted. The population vector cross-correlation was calculated similarly by randomly drawing half of the trials to construct one vector and using the other 50% trials to construct the other vector. For each spatial bin, the Pearson correlation coefficient was calculated. The location reconstruction error was calculated as the distance between the spatial bin with the highest cross-correlation value to the animals’ actual location. As randomly splitting trials into halves can lead to spurious cross-correlation maps, this process was repeated 100 times and the mean cross-correlation coefficient and position reconstruction error for each spatial bin was calculated.

### Activity during decoupled stimulus presentation

The peak response during decoupled stimulus presentation was evaluated by aligning each neuron or bouton to its preferred alignment point during virtual navigation. The response was then measured at the same point relative to that alignment point (in space) where it showed its peak response during virtual navigation. To allow for small shifts in peak activity during decoupled stimulus presentation, a window of ±20 cm was introduced and the peak value within that window was used for analysis. Transients were identified whenever ΔF/F in a given trial (binned into 2 cm spatial bins) rose above 6 standard deviations of that neurons baseline activity (70^th^ percentile of data points) for at least 2 consecutive bins. Transients located outside ±60 cm of the mean peak during virtual navigation were excluded. Jitter was calculated as the standard error of the difference between mean peak and transient peak locations.

### Motor vs. no motor analysis

The effect of concurrent motor and visual inputs during decoupled stimulus presentation was assessed by grouping trials based on whether the animal was running or stationary. A trial was considered a ‘running’ trial if its average speed exceeded 3 cm/sec in a ±50 cm time window around its peak response relative to the landmark during virtual navigation. To allow for slight mismatches between a neuron’s peak response during virtual navigation and decoupled stimulus presentation, the peak within 20 cm of the VR peak was used. Activity in the black box was calculated as a neuron’s response 1.5 seconds after onset of showing black screens with a movement time window of ±1 second. The relative response amplitudes were calculated by normalizing the activity of each neuron to its activity during virtual navigation. Only sessions in which a given animal ran and was stationary were included in this analysis.

### Laminar analysis

To assess differences in neuronal responses of superficial vs deep neurons in RSC, the depth of the recordings was used to determine which layer neurons belonged to. RSC does not possess a layer 4, and layers 2/3 and 5 are separated by a section of relatively few cell bodies. In addition, layer 5 is comparatively superficial, starting at only 300 µm below the pia (Lein et al., 2007). This made identification of cortical layers during in vivo 2-photon imaging possible. We split recordings into layer 2/3 and layer 5 recordings based on depth below the pia. In one recording, a Rbp4-Cre positive animal, which expresses Cre in many layer 5 cells, was used in conjunction with a flexed GCaMP6f construct. The recording depth for this animal was congruent with other recordings in which we located layer 5 based on recording depth alone.

## Acknowledgements

We are grateful to Ben Scott, Jeff Gauthier, Elias Issa, Matt Wilson, and Ashok Litwin-Kumar for comments on the manuscript. We thank Enrique Toloza, Jakob Voigts, and the rest of the Harnett lab for critical feedback on the project, analyses, and manuscript. Support was provided by NIH RO1NS106031, the McGovern Institute for Brain Research, the NEC Corporation Fund for Research in Computers and Communications at MIT, and the Klingenstein-Simons Fellowship in Neuroscience (to M.T.H.).

## Contributions

L.F.F. designed the experiments, performed surgeries and behavioral training, conducted behavioral and imaging experiments, analyzed the data, prepared the figures, and co-wrote the manuscript. R.M.S-A assisted with training, behavioral and imaging experiments, data acquisition, and data analysis. F.B. helped design the behavioral paradigm and assisted with training and analysis. M.T.H conceived, supervised the project and co-wrote the manuscript.

## Supplemental Information

**Supplemental Figure 1:**
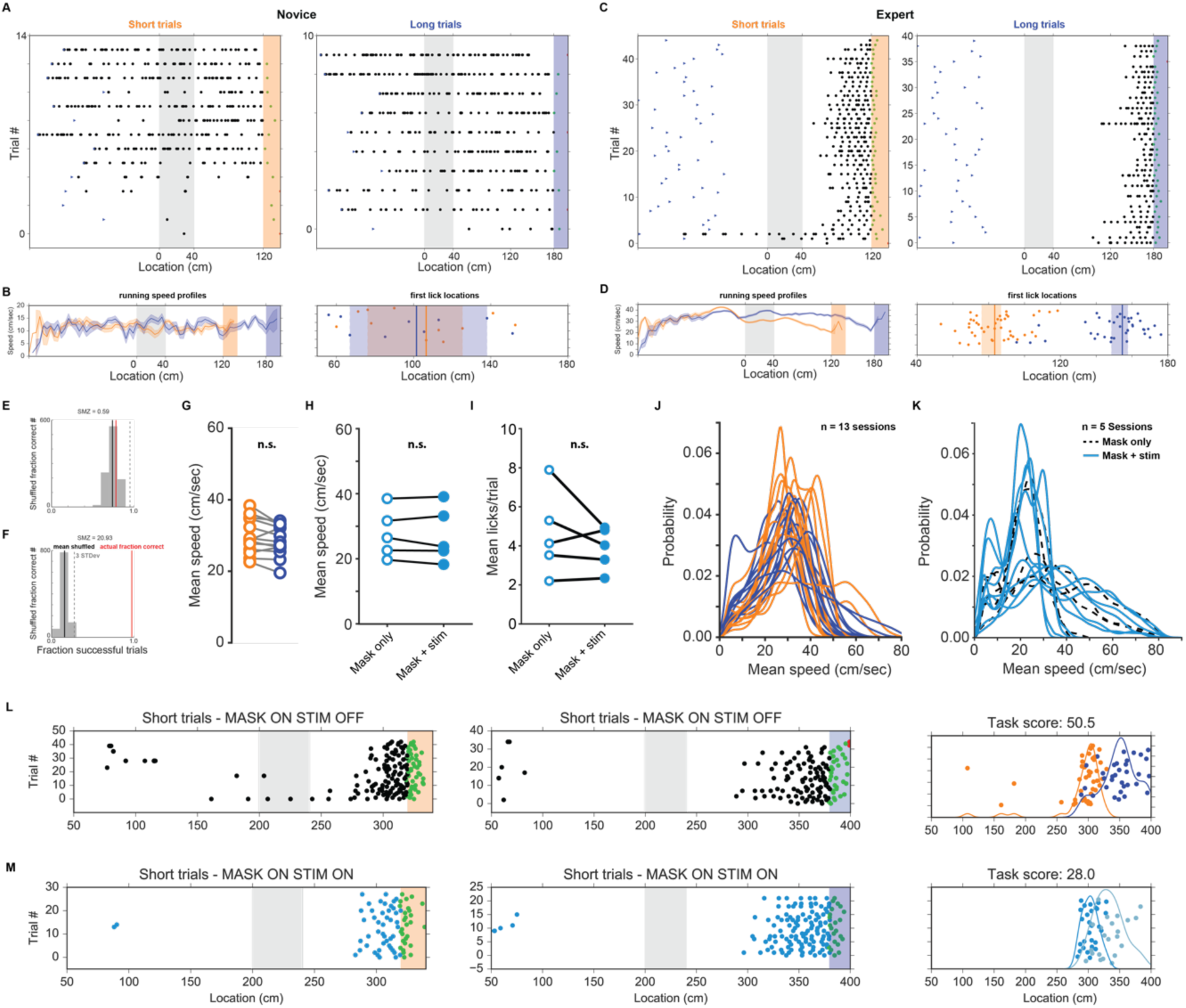
Running and licking behavior. (**A**) Raster plot of licking behavior of example session of a novice animal with short and long trials separated. Note: during the recording the trials were interleaved. The blue triangles indicate the start location of each trial. (**B**) Left: Mean running speed as a function of space (5 cm bins, shaded is standard error of the mean). Right: Location of first licks on short and long trials superimposed. Shaded areas indicate 95% confidence interval. (**C, D**) Same as A & B but for an expert animal. (**E**) Spatial modulation z-score (SMZ) for novice animal (shown in (A)). The grey bars represent a histogram of fraction of successful trials when the location of licks was rotated randomly (repeated 1000 times). Dashed line: 3 standard deviations of that distribution. The red line indicated the animal’s actual fraction of successful trials within that session. (**F**) SMZ of an expert animal (shown in (B)). (**G**) Mean running speed on short and long trials for all recording sessions (Wilcoxon signed-rank test: p > 0.05) (**H**) Same analysis as (D) but for running speed during mask only and mask + stimulation trials during optogenetic inactivation sessions. (**I**) Mean number of licks in mask only vs mask + stim conditions. Overall number of licks was not influenced by stimulation light. (**J**) Kernel density estimates of running speeds for all recording sessions for short and long trials. (**K**) Same analysis as (J) but during mask only and mask + stim conditions. (**L**) Licking behavior in a session where on 50% of trials only the masking light was shown, and the other 50% the masking and optogenetic stimulation light was shown. On short trials (left column) and long trials (center column) when only the masking light was shown. Right column: the first lick per trial on short (orange) and long (blue) trials plotted. (**M**) Same as (L), but when mask and optogenetic stimulation light was on.

**Supplemental Figure 2:**
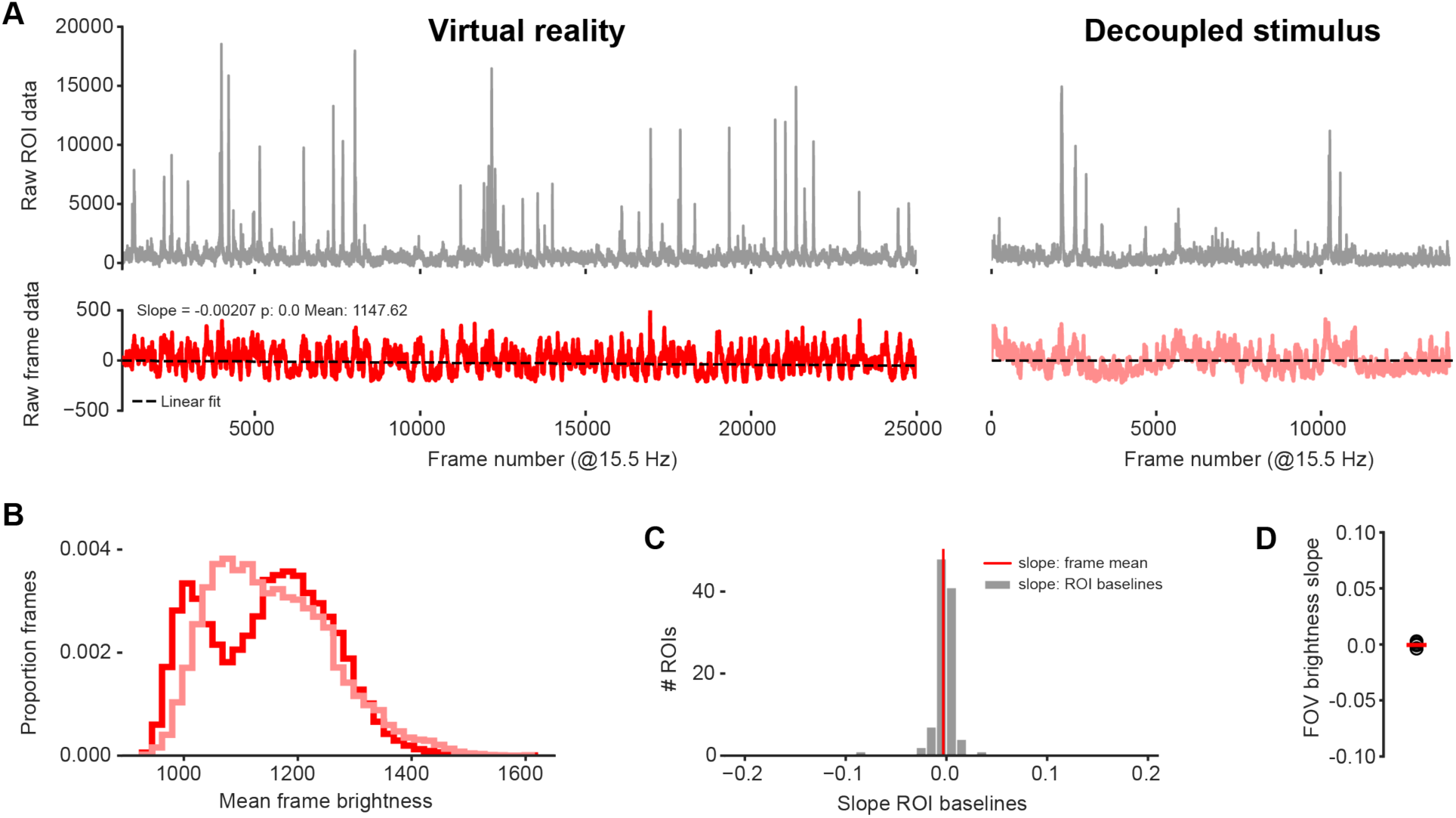
Stable long-term imaging. (**A**) Raw brightness data from one example ROI (gray, top) and whole field-of-view (FOV) brightness (red, bottom) during virtual navigation (left) and subsequent decoupled stimulus presentation (right). Dashed lines across red traces are linear regression fits. (**B**) Mean frame brightness of all frames during virtual navigation and decoupled stimulus presentation. (**C**) Regression line slopes of all ROIs in FOV from which examples are shown in (A) and (B). (**D**) Regression line slopes of FOVs for all recording sessions of RSC neurons contained in this study.

**Supplemental Figure 3:**
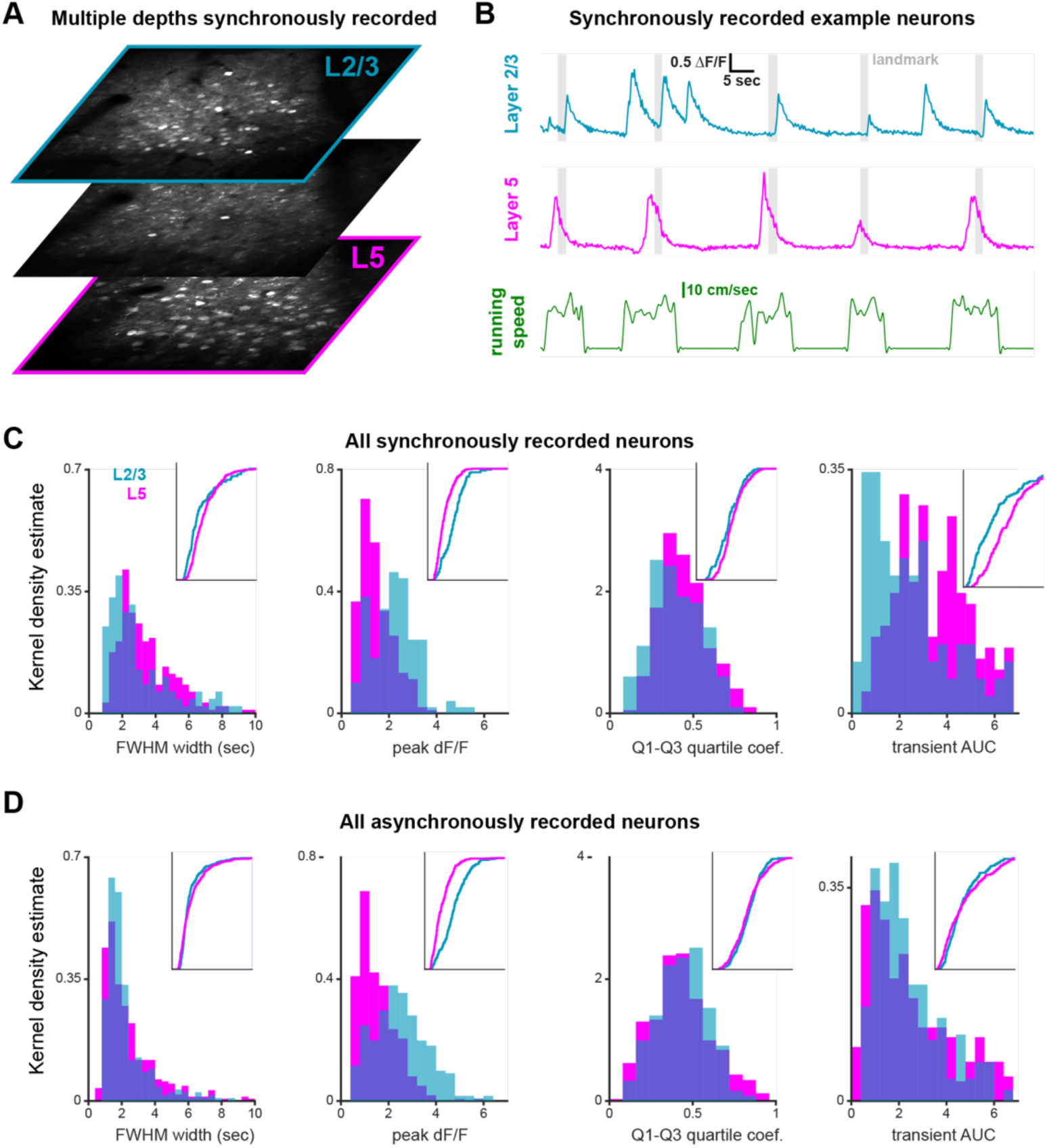
Simultaneous imaging of layer 2/3 and 5. (**A**) Mean images of layers 2/3, 5, and an intermediate imaging plane (3 out of a total of 6). Note the almost total lack of cell bodies in the intermediate layer. (**B**) Two example traces and the corresponding running speed. (**C**) Properties of transients of all simultaneously recorded neurons in L2/3 and L5 neurons (n = 3 sessions from 3 mice. Note: only one session is included in data of main figures as behavior was below threshold for the other two.). Full width at half maximum (FWHM), Q1-Q3 quartile coefficient, and area under the curve (AUC) were broadly similar. Only peak ΔF/F differed between layers, which is likely explained by differences in optical access to superficial vs deep layers. (**D**) Same data as in (C) but this time from asynchronously recorded neurons (n = 7 sessions from 5 animals, all included in main figures). Properties for synchronously and asynchronously recorded neurons are broadly similar; only AUC differs, which may be explained by the impact stray fluorescence from overlaying GCaMP6f expression neurons.

**Supplemental Figure 4:**
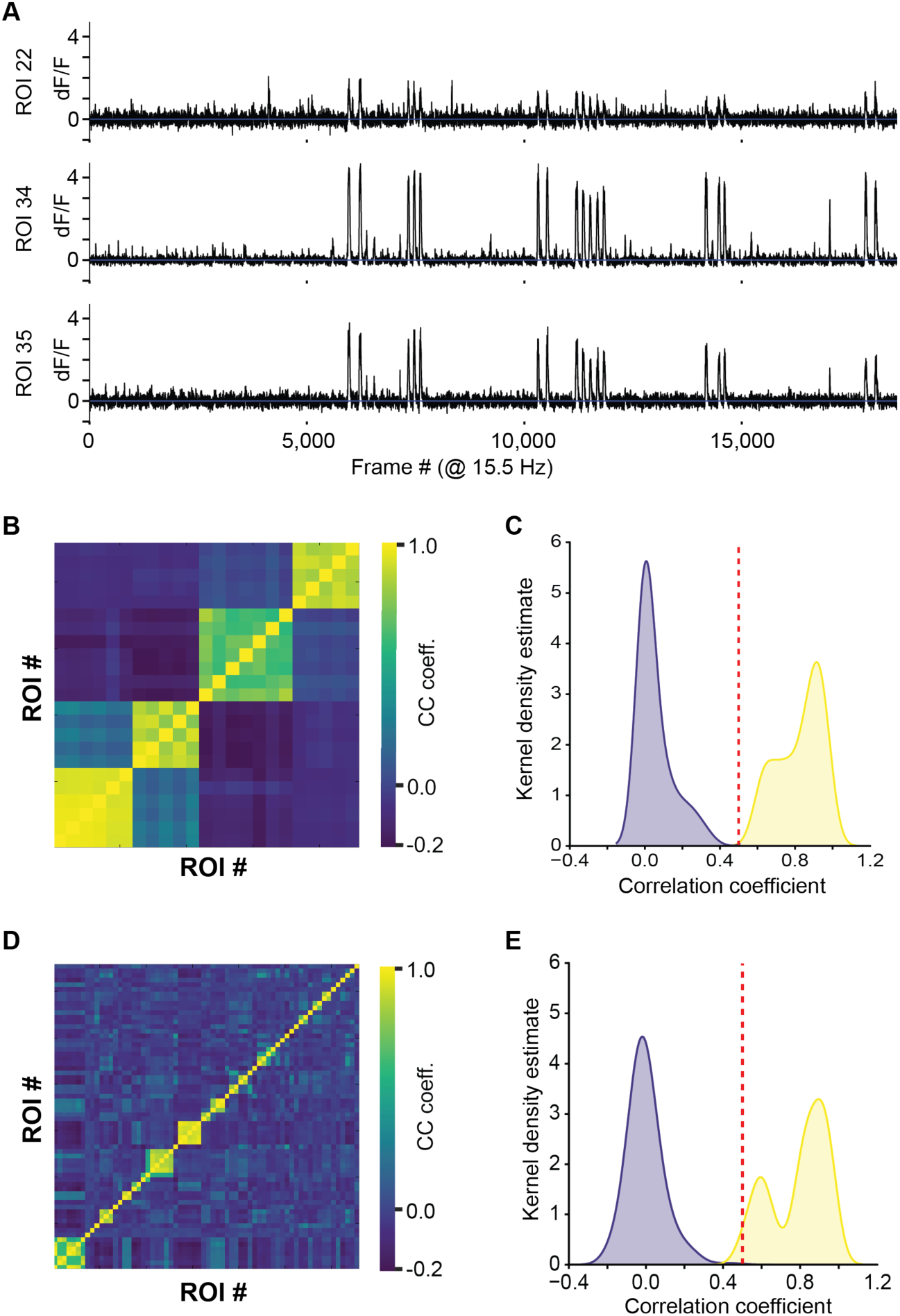
Identification process for unique axonal inputs. (**A**) Traces of three bouton ROIs, putatively from the same axon. (**B**) Cross-correlation matrix of 23 boutons, manually identified to belong to 4 individual axons. (**C**) Cross-correlation of all boutons in (B), split into those belonging to the same axon (yellow) and different axons (blue). The threshold was set manually at 0.5 based on this distribution. (**D**) The cross-correlation matrix was ordered by grouping together all ROIs above the threshold estimated in (C). (**E**) Same plot as (C). Confirmation that the threshold in (C) is valid for the whole population.

**Supplemental Figure 5:**
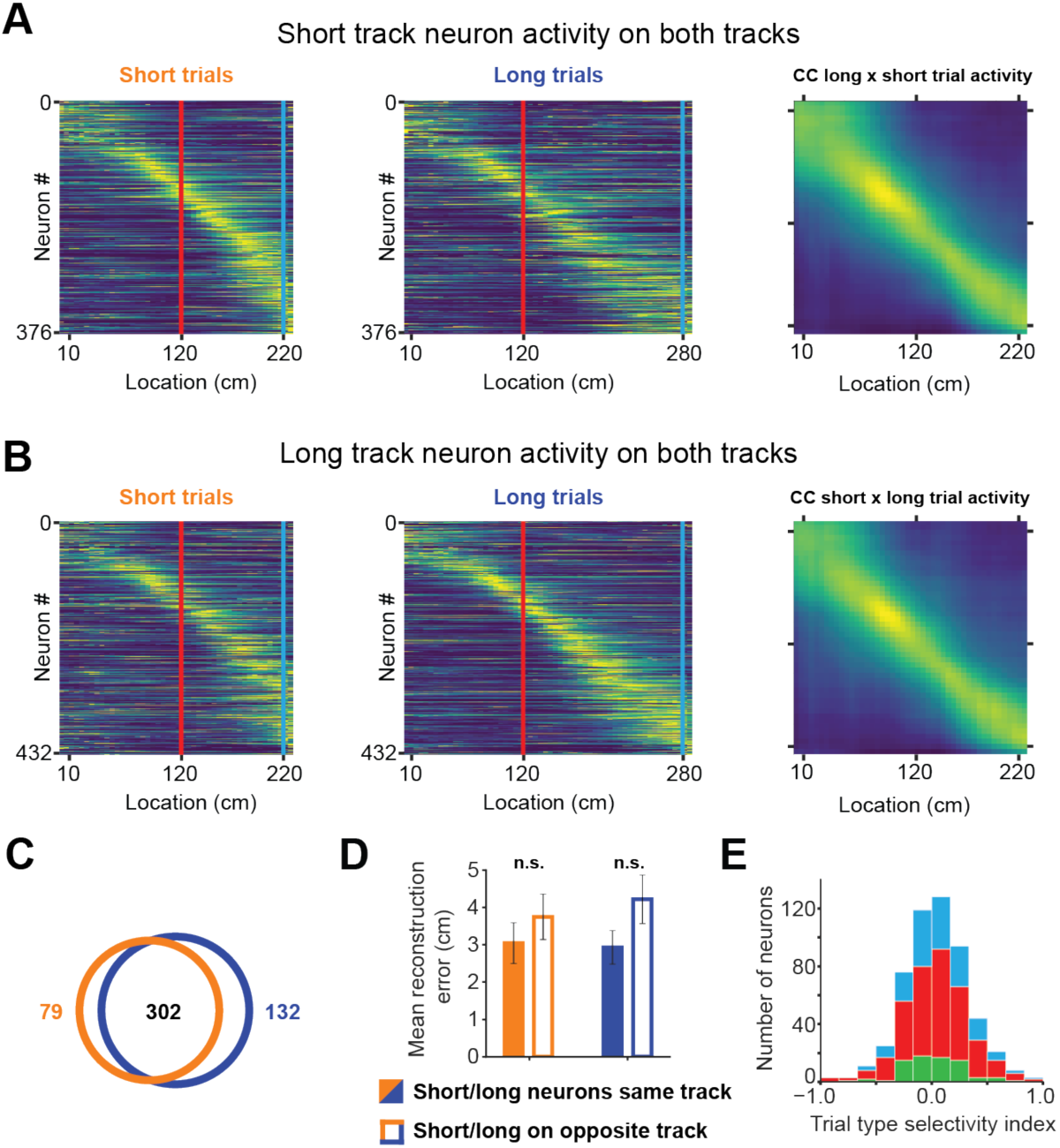
Short track and long track neurons activity on opposite tracks. (**A**) Activity of short track neurons on the short track (left), long track (middle) and the cross correlation of population vectors on each track (right). (**B**) Same as (A) but with long track neuron population. (**C**) Venn diagram showing the overlap of neural populations. The majority of neurons is active on both tracks with only a small fraction being active on exclusively one or the other. (**D**) Location reconstruction error using only short track or long track neurons on the respective opposite track is slightly, but not significantly larger compared to the same track (mean reconstruction error_short/short_: 3.04 ± 0.55, error_short/long_: 3.75 ± 3.75; error _long/long_: 2.93 ± 0.44, error _long/short_: 4.22 ± 0.65; unpaired t-test short track active neurons: p = 0.39, long track active neurons: p = 0.1). (**E**) Stacked bar chart trial type selectivity index [(Response_short_ – Response_long_)/(Response_short_ + Response_long_)] for all three cell types.

## Bibliography

Alexander, A. S., & Nitz, D. A. (2015). Retrosplenial cortex maps the conjunction of internal and external spaces. Nature Neuroscience, 18(8), 1143–1151. https://doi.org/10.1038/nn.4058

Alexander, A. S., & Nitz, D. A. (2017). Spatially Periodic Activation Patterns of Retrosplenial Cortex Encode Route Sub-spaces and Distance Traveled. Current Biology, 27(11), 1551–1560.e4. https://doi.org/10.1016/j.cub.2017.04.036

Aronov, D., & Tank, D. W. (2014). Engagement of Neural Circuits Underlying 2D Spatial Navigation in a Rodent Virtual Reality System. Neuron, 84(2), 442–456. https://doi.org/10.1016/j.neuron.2014.08.042

Attinger, A., Wang, B., & Keller, G. B. (2017). Visuomotor Coupling Shapes the Functional Development of Mouse Visual Cortex. Cell, 169(7), 1291–1302.e14. https://doi.org/10.1016/j.cell.2017.05.023

Auger, S. D., Mullally, S. L., & Maguire, E. A. (2012). Retrosplenial cortex codes for permanent landmarks. PLoS ONE, 7(8). https://doi.org/10.1371/journal.pone.0043620

Bittner, K. C., Grienberger, C., Vaidya, S. P., Milstein, A. D., Macklin, J. J., Suh, J., … Magee, J. C. (2015). Conjunctive input processing drives feature selectivity in hippocampal CA1 neurons. Nature Neuroscience, 18(8), 1133–1142. https://doi.org/10.1038/nn.4062

Burak, Y., & Fiete, I. R. (2009). Accurate path integration in continuous attractor network models of grid cells. PLoS Computational Biology, 5(2), e1000291. https://doi.org/10.1371/journal.pcbi.1000291

Burgess, N., Barry, C., & O’Keefe, J. (2007). An oscillatory interference model of grid cell firing. Hippocampus, 17(9), 801–812. https://doi.org/10.1002/hipo.20327

Buzsáki, G. (2005). Theta rhythm of navigation: link between path integration and landmark navigation, episodic and semantic memory. Hippocampus, 15(7), 827–840. https://doi.org/10.1002/hipo.20113

Campbell, M. G., Ocko, S. A., Mallory, C. S., Low, I. I. C., Ganguli, S., & Giocomo, L. M. (2018). Principles governing the integration of landmark and self-motion cues in entorhinal cortical codes for navigation. Nature Neuroscience, 21(8), 1096–1106. https://doi.org/10.1038/s41593-018-0189-y

Cho, J., & Sharp, P. E. (2001). Head direction, place, and movement correlates for cells in the rat retrosplenial cortex. Behavioral Neuroscience, 115(1), 3–25. https://doi.org/10.1037/0735-7044.115.1.3

Clancy, K. B., Orsolic, I., & Mrsic-Flogel, T. D. (2019). Locomotion-dependent remapping of distributed cortical networks. Nature Neuroscience, 22(5), 778–786. https://doi.org/10.1038/s41593-019-0357-8

Cooper, B. G., Manka, T. F., & Mizumori, S. J. Y. (2001). Finding your way in the dark: The retrosplenial cortex contributes to spatial memory and navigation without visual cues. Behavioral Neuroscience, 115(5), 1012–1028. https://doi.org/10.1037/0735-7044.115.5.1012

Cooper, B. G., & Mizumori, S. J. Y. (1999). Retrosplenial cortex inactivation selectively impairs navigation in darkness. NeuroReport, 10(3), 625–630. https://doi.org/10.1097/00001756-199902250-00033

Elduayen, C., & Save, E. (2014). The retrosplenial cortex is necessary for path integration in the dark. Behavioural Brain Research, 272, 303–307. https://doi.org/10.1016/j.bbr.2014.07.009

Epstein, R. A. (2008). Parahippocampal and retrosplenial contributions to human spatial navigation. Trends in Cognitive Sciences, 12(10), 388–396. https://doi.org/10.1016/j.tics.2008.07.004

Epstein, R. A., & Vass, L. K. (2014). Neural systems for landmark-based wayfinding in humans. Philosophical Transactions of the Royal Society of London. Series B, Biological Sciences, 369(1635), 20120533. https://doi.org/10.1098/rstb.2012.0533

Etienne, A. S., & Jeffery, K. J. (2004). Path integration in mammals. Hippocampus, 14(2), 180–192. https://doi.org/10.1002/hipo.10173

Etienne, A. S., Maurer, R., & Séguinot, V. (1996). Path integration in mammals and its interaction with visual landmarks. The Journal of Experimental Biology, 199(Pt 1), 201–209. Retrieved from http://www.ncbi.nlm.nih.gov/pubmed/11769306

Fiser, A., Mahringer, D., Oyibo, H. K., Petersen, A. V, Leinweber, M., & Keller, G. B. (2016). Experience-dependent spatial expectations in mouse visual cortex. Nature Neuroscience, (September). https://doi.org/10.1038/nn.4385

Fuhs, M. C., & Touretzky, D. S. (2006). A spin glass model of path integration in rat medial entorhinal cortex. The Journal of Neuroscience : The Official Journal of the Society for Neuroscience, 26(16), 4266–4276. https://doi.org/10.1523/JNEUROSCI.4353-05.2006

Funamizu, A., Kuhn, B., & Doya, K. (2016). Neural substrate of dynamic Bayesian inference in the cerebral cortex. *Nature Neuroscience*, (September). https://doi.org/10.1038/nn.4390

Gauthier, J. L., & Tank, D. W. (2018). A Dedicated Population for Reward Coding in the Hippocampus. Neuron, 99(1), 179–193.e7. https://doi.org/10.1016/j.neuron.2018.06.008

Gothard, K. M., & Skaggs, W. E. (1996). Binding of hippocampal CA1 neural activity to multiple reference frames in a landmark-based navigation task. The Journal of …, 16(2), 823–835. Retrieved from http://psycnet.apa.org/psycinfo/1996-93738-003

Gothard, K. M., Skaggs, W. E., & McNaughton, B. L. (1996). Dynamics of Mismatch Correction in the Hippocampal Ensemble Code for Space: Interaction between Path Integration and Environmental Cues. J. Neurosci., 16(24), 8027–8040. Retrieved from http://www.jneurosci.org.ezproxy.is.ed.ac.uk/content/16/24/8027.long

Groen, T. v, & Wyss, J. M. (1992). Connections of the retrosplenial dysgranular cortex in the rat. The Journal of Comparative Neurology, 315(2), 200–216. https://doi.org/10.1002/cne.903150207

Guo, Z. V, Hires, A. S., Li, N., O’Connor, D. H., Komiyama, T., Ophir, E., … Svoboda, K. (2014). Procedures for behavioral experiments in head-fixed mice. PloS One, 9(2), e88678. https://doi.org/10.1371/journal.pone.0088678

Hafting, T., Fyhn, M., Molden, S., Moser, M., & Moser, E. I. (2005). Microstructure of a spatial map in the entorhinal cortex. Nature, 436(7052), 801–806. https://doi.org/10.1038/nature03721

Hardcastle, K., Ganguli, S., & Giocomo, L. M. (2015). Environmental Boundaries as an Error Correction Mechanism for Grid Cells. Neuron, 86(3), 827–839. https://doi.org/10.1016/j.neuron.2015.03.039

Harnett, M. T., Magee, J. C., & Williams, S. R. (2015). Distribution and Function of HCN Channels in the Apical Dendritic Tuft of Neocortical Pyramidal Neurons. Journal of Neuroscience, 35(3), 1024–1037. https://doi.org/10.1523/JNEUROSCI.2813-14.2015

Harvey, C. D., Coen, P., & Tank, D. W. (2012). Choice-specific sequences in parietal cortex during a virtual-navigation decision task. Nature, 484(7392), 62–68. https://doi.org/10.1038/nature10918

Harvey, C. D., Collman, F., Dombeck, D. A., & Tank, D. W. (2009). Intracellular dynamics of hippocampal place cells during virtual navigation. Nature, 461(7266), 941–946. https://doi.org/10.1038/nature08499

Ino, T., Doi, T., Hirose, S., Kimura, T., Ito, J., & Fukuyama, H. (2007). Directional disorientation following left retrosplenial hemorrhage: A case report with fMRI studies. Cortex, 43(2), 248–254. https://doi.org/10.1016/S0010-9452(08)70479-9

Jacob, P., Casali, G., Spieser, L., Page, H., Overington, D., & Jeffery, K. J. (2017). An independent, landmark-dominated head-direction signal in dysgranular retrosplenial cortex. Nature Neuroscience, 20(2), 173–175. https://doi.org/10.1038/nn.4465

Jadi, M. P., Behabadi, B. F., Poleg-Polsky, A., Schiller, J., & Mel, B. W. (2014). An Augmented Two-Layer Model Captures Nonlinear Analog Spatial Integration Effects in Pyramidal Neuron Dendrites. Proceedings of the IEEE, 102(5), 782–798. https://doi.org/10.1109/JPROC.2014.2312671

Jeffery, K. J. (1998). Learning of landmark stability and instability by hippocampal place cells. Neuropharmacology, 37(4–5), 677–687. Retrieved from http://www.ncbi.nlm.nih.gov/pubmed/9705005

Ji, D., & Wilson, M. A. (2007). Coordinated memory replay in the visual cortex and hippocampus during sleep. Nature Neuroscience, 10(1), 100–107. https://doi.org/10.1038/nn1825

Ji, N., Magee, J. C., & Betzig, E. (2008). High-speed, low-photodamage nonlinear imaging using passive pulse splitters. Nature Methods, 5(2), 197–202. https://doi.org/10.1038/nmeth.1175

Jones, E., Oliphant, T., & Peterson, P. (2001). SciPy: Open source scientific tools for Python. Retrieved from http://www.scipy.org/

Julian, J. B., Keinath, A. T., Marchette, S. A., & Epstein, R. A. (2018). The Neurocognitive Basis of Spatial Reorientation. Current Biology, 28(17), R1059–R1073. https://doi.org/10.1016/j.cub.2018.04.057

Knierim, J. J., Kudrimoti, H., & McNaughton, B. L. (1995). Place cells, head direction cells, and the learning of landmark stability. The Journal of …, 15(March), 1648–1659. Retrieved from http://www.jneurosci.org/content/15/3/1648.short

Knierim, J. J., Kudrimoti, H., & McNaughton, B. L. (1998). Interactions between idiothetic cues and external landmarks in the control of place cells and head direction cells. Journal of …, 425–446. Retrieved from http://krieger.jhu.edu/mbi/knierimlab/publications/Ideotheticcuesandexternallandmarks.pdf

Koay, S. A., Thiberge, S. Y., Brody, C. D., & Tank, D. W. (2019). Neural Correlates of Cognition in Primary Visual versus Neighboring Posterior Cortices during Visual Evidence-Accumulation-based Navigation. Bioarxiv, 1–84. https://doi.org/10.1101/568766

Kononenko, N. L., & Witter, M. P. (2012). Presubiculum layer III conveys retrosplenial input to the medial entorhinal cortex. Hippocampus, 22(4), 881–895. https://doi.org/10.1002/hipo.20949

Larkum, M. E., Kaiser, K. M., & Sakmann, B. (1999). Calcium electrogenesis in distal apical dendrites of layer 5 pyramidal cells at a critical frequency of back-propagating action potentials. Proceedings of the National Academy of Sciences of the United States of America, 96(25), 14600–14604. Retrieved from http://www.ncbi.nlm.nih.gov/pubmed/10588751%0Ahttp://www.pubmedcentral.nih.gov/articlerender.fcgi?artid=PMC24482

Lein, E. S., Hawrylycz, M. J., Ao, N., Ayres, M., Bensinger, A., Bernard, A., … Jones, A. R. (2007). Genome-wide atlas of gene expression in the adult mouse brain. Nature, 445(7124), 168–176. https://doi.org/10.1038/nature05453

Lewis, L. D., Voigts, J., Flores, F. J., Ian Schmitt, L., Wilson, M. A., Halassa, M. M., & Brown, E. N. (2015). Thalamic reticular nucleus induces fast and local modulation of arousal state. ELife, 4(OCTOBER2015), 1–23. https://doi.org/10.7554/eLife.08760

Liu, D., Gu, X., Zhu, J., Zhang, X., Han, Z., Yan, W., … Li, C. T. (2014). Medial prefrontal activity during delay period contributes to learning of a working memory task. Science, 346(6208), 458–463. https://doi.org/10.1126/science.1256573

London, M., & Häusser, M. (2005). Dendritic computation. Annual Review of Neuroscience, 28, 503–532. https://doi.org/10.1146/annurev.neuro.28.061604.135703

Maguire, E. A. (2001). The retrosplenial contribution to human navigation: a review of lesion and neuroimaging findings. Scandinavian Journal of Psychology, 42(3), 225–238. https://doi.org/10.1111/1467-9450.00233

Makino, H., & Komiyama, T. (2015). Learning enhances the relative impact of top-down processing in the visual cortex. Nature Neuroscience, 18(8), 1116–1122. https://doi.org/10.1038/nn.4061

Mante, V., Sussillo, D., Shenoy, K. V., & Newsome, W. T. (2013). Context-dependent computation by recurrent dynamics in prefrontal cortex. Nature, 503(7474), 78–84. https://doi.org/10.1038/nature12742

Mao, D., Kandler, S., McNaughton, B. L., & Bonin, V. (2017). Sparse orthogonal population representation of spatial context in the retrosplenial cortex. Nature Communications, 8(1), 243. https://doi.org/10.1038/s41467-017-00180-9

McNaughton, B. L., Battaglia, F. P., Jensen, O., Moser, E. I., & Moser, M. (2006). Path integration and the neural basis of the “cognitive map”. Nature Reviews. Neuroscience, 7(8), 663–678. https://doi.org/10.1038/nrn1932

McNaughton, B. L., Chen, L. L., & Markus, E. J. (1991). “Dead reckoning,” landmark learning, and the sense of direction: a neurophysiological and computational hypothesis. Journal of Cognitive Neuroscience, 3(2), 190–202. https://doi.org/10.1162/jocn.1991.3.2.190

Miller, E. K., & Cohen, J. D. (2001). An Integrative Theory of Prefrontal Cortex Function. Annual Review of Neuroscience, 24(1), 167–202. https://doi.org/10.1146/annurev.neuro.24.1.167

Minoshima, S., Giordani, B., Berent, S., Frey, K. A., Foster, N. L., & Kuhl, D. E. (1997). Metabolic reduction in the posterior cingulate cortex in very early Alzheimer’s disease. Annals of Neurology, 42(1), 85–94. https://doi.org/10.1002/ana.410420114

Miyashita, T., & Rockland, K. S. (2007). GABAergic projections from the hippocampus to the retrosplenial cortex in the rat. The European Journal of Neuroscience, 26(5), 1193–1204. https://doi.org/10.1111/j.1460-9568.2007.05745.x

Monaco, J. D., Knierim, J. J., & Zhang, K. (2011). Sensory Feedback, Error Correction, and Remapping in a Multiple Oscillator Model of Place-Cell Activity. Frontiers in Computational Neuroscience, 5(September), 1–17. https://doi.org/10.3389/fncom.2011.00039

Montijn, J. S., Vinck, M., & Pennartz, C. M. A. (2014). Population coding in mouse visual cortex: response reliability and dissociability of stimulus tuning and noise correlation. Frontiers in Computational Neuroscience, 8(June), 1–15. https://doi.org/10.3389/fncom.2014.00058

Morris, R. G. M. (1981). Spatial localization does not require the presence of local cues. Learning and Motivation, 12(2), 239–260. https://doi.org/10.1016/0023-9690(81)90020-5

Muller, R. U., & Kubie, J. L. (1987). The effects of changes in the environment on the spatial firing of hippocampal complex-spike cells. J Neurosci, 7(July). Retrieved from http://www.researchgate.net/publication/19550157_The_effects_of_changes_in_the_environment_on_the_spatial_firing_of_hippocampal_complex-spike_cells/file/d912f50c0b9519ca49.pdf

Murray, J. D., Bernacchia, A., Roy, N. A., Constantinidis, C., Romo, R., & Wang, X. (2017). Stable population coding for working memory coexists with heterogeneous neural dynamics in prefrontal cortex. Proceedings of the National Academy of Sciences, 114(2), 394–399. https://doi.org/10.1073/pnas.1619449114

Newman, J. P., Fong, M., Millard, D. C., Whitmire, C. J., Stanley, G. B., & Potter, S. M. (2015). Optogenetic feedback control of neural activity. ELife, 4(JULY 2015), 1–24. https://doi.org/10.7554/eLife.07192

Niell, C., & Stryker, M. P. (2010). Modulation of visual responses by behavioral state in mouse visual cortex. Neuron, 65(4), 472–479. https://doi.org/10.1016/j.neuron.2010.01.033

Oh, S. W., Harris, J. A., Ng, L., Winslow, B., Cain, N., Mihalas, S., … Zeng, H. (2014). A mesoscale connectome of the mouse brain. Nature, 508(7495), 207–214. https://doi.org/10.1038/nature13186

Pakan, J. M., Currie, S. P., Fischer, L., & Rochefort, N. L. (2018). The Impact of Visual Cues, Reward, and Motor Feedback on the Representation of Behaviorally Relevant Spatial Locations in Primary Visual Cortex. Cell Reports, 24(10), 2521–2528. https://doi.org/10.1016/j.celrep.2018.08.010

Pengas, G., Hodges, J. R., Watson, P., & Nestor, P. J. (2010). Focal posterior cingulate atrophy in incipient Alzheimer’s disease. Neurobiology of Aging, 31(1), 25–33. https://doi.org/10.1016/j.neurobiolaging.2008.03.014

Pérez-Escobar, J. A., Kornienko, O., Latuske, P., Kohler, L., & Allen, K. (2016). Visual landmarks sharpen grid cell metric and confer context specificity to neurons of the medial entorhinal cortex. ELife, 5(JULY), 1232627. https://doi.org/10.7554/eLife.16937

Poort, J., Khan, A. G., Pachitariu, M., Nemri, A., Orsolic, I., Krupic, J., … Hofer, S. B. (2015). Learning Enhances Sensory and Multiple Non-sensory Representations in Primary Visual Cortex. Neuron. The Authors. https://doi.org/10.1016/j.neuron.2015.05.037

Ranganathan, G. N., Apostolides, P. F., Harnett, M. T., Xu, N. L., Druckmann, S., & Magee, J. C. (2018). Active dendritic integration and mixed neocortical network representations during an adaptive sensing behavior. Nature Neuroscience, 21(11), 1583–1590. https://doi.org/10.1038/s41593-018-0254-6

Rigotti, M., Barak, O., Warden, M. R., Wang, X., Daw, N. D., Miller, E. K., & Fusi, S. (2013). The importance of mixed selectivity in complex cognitive tasks. Nature, 497(7451), 585–590. https://doi.org/10.1038/nature12160

Robertson, C. E., Hermann, K. L., Mynick, A., Kravitz, D. J., & Kanwisher, N. (2016). Neural Representations Integrate the Current Field of View with the Remembered 360° Panorama in Scene-Selective Cortex. Current Biology, 26(18), 2463–2468. https://doi.org/10.1016/j.cub.2016.07.002

Saleem, A. B., Diamanti, M. E., Fournier, J., Harris, K. D., & Carandini, M. (2018). Coherent encoding of subjective spatial position in visual cortex and hippocampus. Nature, 562(7725), 124–127. https://doi.org/10.1038/s41586-018-0516-1

Smith, D. M., Barredo, J., & Mizumori, S. J. Y. (2012). Complimentary roles of the hippocampus and retrosplenial cortex in behavioral context discrimination. Hippocampus, 22(5), 1121–1133. https://doi.org/10.1002/hipo.20958

Spiers, H. J., & Maguire, E. A. (2006). Thoughts, behaviour, and brain dynamics during navigation in the real world. NeuroImage, 31, 1826–1840. https://doi.org/10.1016/j.neuroimage.2006.01.037

Sreenivasan, S., & Fiete, I. R. (2011). Grid cells generate an analog error-correcting code for singularly precise neural computation. Nature Neuroscience, 14(10), 1330–1337. https://doi.org/10.1038/nn.2901

Stringer, C., Pachitariu, M., Steinmetz, N., Carandini, M., & Harris, K. D. (2019). High-dimensional geometry of population responses in visual cortex. Nature. https://doi.org/10.1038/s41586-019-1346-5

Sugar, J., Witter, M. P., van Strien, N. M., & Cappaert, N. L. M. (2011). The retrosplenial cortex: intrinsic connectivity and connections with the (para)hippocampal region in the rat. An interactive connectome. Frontiers in Neuroinformatics, 5, 7. https://doi.org/10.3389/fninf.2011.00007

Svoboda, E., McKinnon, M. C., & Levine, B. (2006). The functional neuroanatomy of autobiographical memory: A meta-analysis. Neuropsychologia, 44(12), 2189–2208. https://doi.org/10.1016/j.neuropsychologia.2006.05.023

Takahashi, N., Oertner, T. G., Hegemann, P., & Larkum, M. E. (2016). Active cortical dendrites modulate perception. Science, 354(6319), 1587–1590. https://doi.org/10.1126/science.aah6066

Taube, J. S. (2007). The head direction signal: origins and sensory-motor integration. Annual Review of Neuroscience, 30, 181–207. https://doi.org/10.1146/annurev.neuro.29.051605.112854

Taube, J. S., Muller, R. U., & Ranck, J. B. (1990a). Head-direction cells recorded from the postsubiculum in freely moving rats. II. Effects of environmental manipulations. The Journal of Neuroscience, 70(February). Retrieved from http://www.jneurosci.org/content/10/2/436.short

Taube, J. S., Muller, R. U., & Ranck, J. B. (1990b). Head-direction cells recorded from the postsubiculum in freely moving rats. II. Effects of environmental manipulations. The Journal of Neuroscience, 70(February). Retrieved from http://www.jneurosci.org/content/10/2/436.short

Tolman, E. C. (1948). Cognitive maps in rats and men. Psychological Review, 55(4), 189–208. Retrieved from http://www.ncbi.nlm.nih.gov/pubmed/9230435

Valenstein, E., Bowers, D., Verfaellie, M., Heilman, K. M., Day, A., & Watson, R. T. (1987). Retrosplenial amnesia. Brain : A Journal of Neurology, 110 (Pt 6), 1631–1646. Retrieved from http://www.ncbi.nlm.nih.gov/pubmed/3427404

Valerio, S., & Taube, J. S. (2012). Path integration: How the head direction signal maintains and corrects spatial orientation. Nature Neuroscience, 15(10), 1445–1453. https://doi.org/10.1038/nn.3215

Vann, S. D., Aggleton, J. P., & Maguire, E. A. (2009). What does the retrosplenial cortex do? Nature Reviews. Neuroscience, 10(11), 792–802. https://doi.org/10.1038/nrn2733

Vedder, L. C., Miller, A. M. P., Harrison, M. B., & Smith, D. M. (2016). Retrosplenial Cortical Neurons Encode Navigational Cues, Trajectories and Reward Locations During Goal Directed Navigation. Cerebral Cortex, 1–11. https://doi.org/10.1093/cercor/bhw192

Villain, N., Desgranges, B., Viader, F., de la Sayette, V., Mezenge, F., Landeau, B., … Chetelat, G. (2008). Relationships between Hippocampal Atrophy, White Matter Disruption, and Gray Matter Hypometabolism in Alzheimer’s Disease. Journal of Neuroscience, 28(24), 6174–6181. https://doi.org/10.1523/jneurosci.1392-08.2008

Vogt, B. A., & Miller, M. W. (1983). Cortical connections between rat cingulate cortex and visual, motor, and postsubicular cortices. The Journal of Comparative Neurology, 216(2), 192–210. https://doi.org/10.1002/cne.902160207

Xu, N. L., Harnett, M. T., Williams, S. R., Huber, D., O’Connor, D. H., Svoboda, K., & Magee, J. C. (2012). Nonlinear dendritic integration of sensory and motor input during an active sensing task. Nature, 492(7428), 247–251. https://doi.org/10.1038/nature11601

Yoder, R. M., Clark, B. J., Brown, J. E., Lamia, M. V, Shinder, M. E., Taube, J. S., … Valerio, S. (2011). Both visual and idiothetic cues contribute to head direction cell stability during navigation along complex routes. Journal of Neurophysiology, 105(6), 2989–3001. https://doi.org/10.1152/jn.01041.2010

Zhang, S., Xu, M., Kamigaki, T., Hoang Do, J. P., Chang, W. C., Jenvay, S., … Dan, Y. (2014). Selective attention. Long-range and local circuits for top-down modulation of visual cortex processing. Science (New York, N.Y.), 345(6197), 660–665. https://doi.org/10.1126/science.1254126

